# A canonical neural circuit for motion direction estimation

**DOI:** 10.64898/2026.10.06.757049

**Authors:** María Berraquero, Ignacio Serrano-Pedraza, Marcelo Bertalmío

**Affiliations:** Spanish National Research Council (CSIC), Spain; Universidad Politécnica de Madrid, Spain; Universidad Complutense de Madrid, Spain

## Abstract

Abundant experimental data support the view that motion detection is a canonical neural operation, common to a wide variety of species and different stages in the visual stream. However, existing motion models lack this generality, as their predictive power tends to be restricted to some specific visual phenomena, brain regions, or animal taxa. Here, we introduce a neural circuit model that only uses cells selective to cardinal directions, with elements that can be interpreted as emulating some dendritic processes, and we show how it is able to predict a wide range of visual motion data, from basic properties to hitherto unexplained phenomena, surpassing state-of-the-art models implemented as deep neural networks with millions of parameters. The direction-selective cells that our model relies on are present in the visual systems of many classes of animals, from insects to primates, and in visual pathway structures going from the retina to the cortex, suggesting that the proposed model could serve as a canonical neural circuit for estimating motion direction.

## Introduction

Motion vision is a key trait for the survival of sighted animals because of its role in many essential tasks, such as navigation, visual course control, hunting prey, evading predators, and detecting mating partners (Borst and Euler, 2011).

Among visual systems of different species, there are striking similarities, as already pointed out over a century ago by Ramón y Cajal (Ramón y Cajal and Sánchez, 1915). From insects to primates, the first motion computation is direction (Yang and Clandinin, 2018): direction-selective (DS) neurons underlie motion perception (Pasternak and Tadin, 2020), they have been studied for many decades, and they are present in a wide variety of animal species and visual system areas. Technological advances in the past twenty years have shown that the architecture for motion detection is shared by insects and vertebrates (Agrochao et al., 2020), and the analogies in visual circuitry are both retinal and post-retinal (Tanaka and Portugues, 2025). There is ample evidence supporting the theory that natural selection tends to favor a relatively limited range of mathematical solutions for motion detection (Clark and Demb, 2016), and that visual motion detection is a canonical neural computation (Clark and Fitzgerald, 2024), with the abundance of analogies explained by constraints from visual ecology (e.g. light usually comes from above, optical flow patterns from forward motion are similar, etc.) and from the evolutionary conservation of neural function, neural development, and photoreceptor molecular machinery across bilaterally symmetric organisms (Tanaka and Portugues, 2025). For instance, responses to moving plaid stimuli have been reported in insects (Saleem et al., 2012), non-primate vertebrates (Matteucci et al., 2021), non-human primates (Guan et al., 2020) and humans (Adelson and Movshon, 1982), and in the retina (DePiero et al., 2024), thalamus (Merabet et al., 1998), superior colliculus (SC) (DePiero et al., 2024), V1 (Van Kemenade et al., 2014), V2 and V3 (Huk and Heeger, 2002), V4 (Li et al., 2025), MT (Movshon et al., 1985) and MST (Khawaja et al., 2013); other examples are the peripheral drift motion illusion, the reverse-Phi and missing-fundamental illusions, and the detection of both first and second-order motion, all of which have been reported in flies, fish, rodents, cats, monkeys, and humans (Agrochao et al., 2020; Tuthill et al., 2011; Orger et al., 2000; Theobald et al., 2008).

However, despite the wide variety of approaches that have been proposed over the years, the existing algorithms for motion direction estimation are not good candidates for a canonical model, as each is affected by one or more important limitations:

1. the model might be tailored to one type of animal, and not be applicable to a very different species (e.g. the motion model of (Simoncelli and Heeger, 1998) works on primates but its elements do not allow for a direct use on insects);
2. the model might be tailored to one brain region, and not work on others (e.g. the model in (Rust et al., 2006) was developed for MT, but it does not work on V1 (Palagina et al., 2017) nor SC (DePiero et al., 2024));
3. the model might be tailored to explain a specific basic phenomenon, but will not explain another (e.g. models for second-order motion do not predict first-order motion (Lu and Sperling, 2001)).

There is a growing consensus that the actual neural circuitry for motion direction estimation is more complex or flexible than that allowed by current models (Currier et al., 2023; DePiero et al., 2024). Despite the amount of progress and the fact that direction selectivity is the best-understood neural computation in the brain (Wei, 2018), the conceptually modest problem of local motion detection is still not fully understood (Mauss et al., 2017; Yang and Clandinin, 2018; Borst and Groschner, 2023). Nonetheless, in the past few years, a number of key elements and properties of DS cells have been established:

- DS cells rely on active dendrites (Borst and Euler, 2011; Svirskis et al., 2015; Mauss et al., 2017; Currier et al., 2023; Kühn et al., 2025; DePiero et al., 2024);
- DS cells require saturation or dynamic nonlinearities (Badwan et al., 2019; Bach and Atala-Gérard, 2020);
- motion estimation with DS cells involves a post-summation stage (Fiscella et al., 2015; Quaia et al., 2022; Tanaka and Portugues, 2025);
- DS cells compute direction *de novo* in the retina, V1, MT, and MST (He et al., 2026) and in SC (Kühn et al., 2025);
- DS cells in insects and vertebrate retina are tuned to cardinal directions (Borst and Euler, 2011), and in vertebrates they have direct pathways to lateral geniculate nucleus (LGN), SC (DePiero et al., 2024), V1 (via LGN) (Sun et al., 2016; Hillier et al., 2017; Wang et al., 2024; Su et al., 2025), V4 (from SC via LGN, skipping V1) (Schmid et al., 2013) and MT (via LGN, skipping V1) (Sincich et al., 2004);
- DS cells tuned to cardinal directions are, as a population, sufficient to properly encode motion (Fiscella et al., 2015).

Considering all the above, in this work, we propose a neural circuit model for the estimation of motion direction that: (1) uses only DS cells tuned to cardinal directions; (2) extends a recent model for 1D motion sensors (Luna et al., 2025) that considers active dendrites and saturating nonlinearities; and (3) includes a post-summation stage. Our experiments show that the proposed model can reproduce a wide variety of results, covering a range of phenomena that go from the basic (contrast saturation, reverse-Phi motion, the missing-fundamental illusion, motion direction estimation for gratings and plaids), through the challenging (motion illusions with static stimuli), and the very challenging (detection of both first and second-order motion, interaction among scales), culminating in the prediction of phenomena that cannot be properly explained with existing models (time-reversal symmetry breaking). We also tested the state-of-the-art in deep neural networks (DNNs) for motion estimation, each of which has millions of parameters and some of which are bio-inspired, and we report that these DNNs are not good vision models as they consistently fail in all the challenging scenarios mentioned above.

Given that our model is based on DS cells tuned to cardinal directions, which are present in very different animal species and in many areas of the visual system, and because our experimental results reproduce a broad scope of motion phenomena observed in very different animals and visual areas, we believe that the proposed model could serve as a canonical neural circuit for estimating motion direction.

## Methods

### Proposed model

Let **I** be the spatio-temporal stimulus, defined over 2D spatial coordinates denoted by *ū* and a temporal interval denoted by 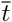. Our proposed model has three stages, each one producing a spatio-temporal output:

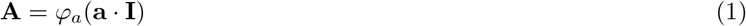

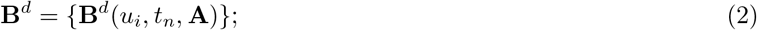

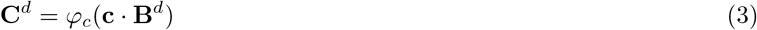

where the first and third stages are spatio-temporal linear filtering operations (**a** and **c**) followed by pointwise nonlinearities (*φ_a_* and *φ_c_*), the superscript *d* denotes the cardinal directions, and the second stage is a modified version of the 1D motion sensor model presented in (Luna et al., 2025), which cannot be written in the compact notation of the other stages due to its highly nonlinear nature (Bertalmío et al., 2020) and for each spatial position *u_i_* and instant *t_n_* is defined as:

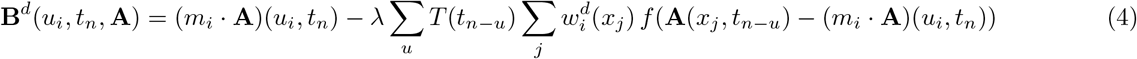

Here *u_i_* = (*x_i_, y_i_*) is a 2D vector with the horizontal and vertical coordinates of point *i, m_i_* and 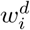 are 2D spatial filters, *λ* is a scalar, *T* is a temporal filter, and *f* (·) is a pointwise nonlinearity. Eq. 4 extends the model of (Luna et al., 2025) by allowing the input arguments of the nonlinearity *f* (·) to include linear and nonlinear transforms of the stimulus.

In a mechanistic interpretation of our model, the linear spatial terms in all three stages would correspond to passive dendrites, and the nonlinear function *f* (·) in the second stage would account for the response of active dendrites, with the subtracting element in its argument representing a back-propagating action potential (bAP).

The output of our model will be the time-varying 2D signals **C***^d^* (Eq. 3), each of them representing the projection of the estimated motion vector into one of the four cardinal directions. For simplicity, we combine the directions in pairs so as to obtain motion-opponent signals corresponding to just two projections, into the horizontal and vertical directions. In other words, *d* ∈ {*H, V*} instead of *d* ∈ {*N, S, E, W*} (which is akin to the recombination carried out in the fly visual system (Clark and Demb, 2016)), and the direction of motion can easily be estimated from these two signals.

### Model instance and parameters

For the experiments, we chose to instantiate our model as follows:

- The function *φ_a_* is the identity and **a** is a filter, Gaussian in space and biphasic in time, that is, band-pass in temporal frequency. The spatial filter *a_s_* has standard deviation *σ_a_*, and the temporal filter follows the equation 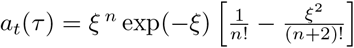 with *n* = 5, where *ξ* = 20 *τ/N_a_* and *τ* denotes the temporal delay over a total duration of *N_a_* both in milliseconds.
- The filter *m_i_* is Gaussian with standard deviation *σ_m_*, and the filters 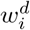 are Gabor functions defined by spatial frequency *f*_0_, envelope width *σ_W_*, and aspect ratio *γ*. The equation for the Gabor filter is: 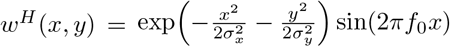 where *σ_x_* = *σ_W_* and *σ_y_* = *σ_W_/γ*. The vertical filter *w^V^* (*x, y*) is obtained by rotating *w^H^* (*x, y*) by 90°.
- Filter *T* is lowpass and is defined as 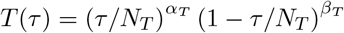 with a total duration of *N_T_* ms.
- The function *f* (·) is a saturating nonlinearity that is positive, even, and zero at the origin:

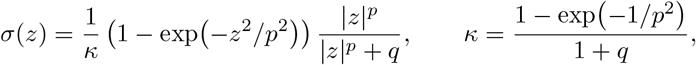

where the same parameter *p* sets both the width of the threshold and the exponent of the saturation term.
- The function *φ_c_* is the identity, and filter **c** is the composition of a spatially variable Gaussian filter in space (*c_s_*) and a monophasic, sustained filter in time (*c_t_*), that is, low-pass in temporal frequency. The width of *c_s_* varies with eccentricity as

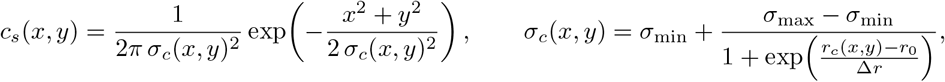

where *r_c_*(*x, y*) is the distance from point (*x, y*) to the image center, *r*_0_ the distance between the image center and the stimulus center, and Δ*r* the width of the transition between *σ*_max_ and *σ*_min_. The temporal filter *c_t_* has the form 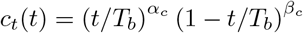 with a total duration of *T_b_*, the duration of the stimulus.

Table 1 indicates the parameter values for each of the experiments.

The estimated motion direction and magnitude at each spatial position and time instant are obtained from the two motion-opponent signals **C***^H^* and **C***^V^* as

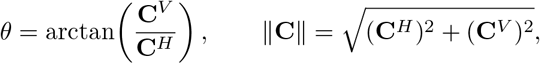

with *θ* measured counterclockwise from the horizontal, so that *θ* = 0 denotes rightward motion.

**Table 1.** Model parameters for each experimental setting.

| Symbol | Unit |  |  |  |  | Breaking time symmetry |  |  | Static motion illusions |  |  |  |  |
| --- | --- | --- | --- | --- | --- | --- | --- | --- | --- | --- | --- | --- | --- |
|  |  | 1D basic phenomena | Second order | Gratings & plaids | Interaction among scales | Opponency (control) | Light edges | Uneven square | Rotating snakes | Central drift | Peripheral drift | Symmetric control | Real rotation |
| $f_0$ | cpd | 2.0 | 2.0 | 2.0 | 1.5 | 2.0 | 2.0 | 2.0 | 1.016 | 4.741 | 2.032 | 1.016 | 1.016 |
| $\sigma_W$ | deg | 0.15 | 0.15 | 0.15 | 0.30 | 0.15 | 0.15 | 0.15 | 0.198 | 0.132 | 0.198 | 0.198 | 0.198 |
| $\gamma$ | — | 0.65 | 0.65 | 0.65 | 0.65 | 0.65 | 0.65 | 0.65 | 0.65 | 0.65 | 0.65 | 0.65 | 0.65 |
| $\sigma_a$ | deg | 0.05 | 0.02 | 0.05 | 0.05 | 0.05 | 0.05 | 0.05 | 0.035 | 0 | 0.035 | 0.035 | 0.035 |
| $\sigma_m$ | deg | 0 | 0 | 0 | 0 | 0 | 0 | 0 | 0 | 0 | 0 | 0 | 0 |
| $\lambda$ | — | 30 | 30 | 30 | 30 | 30 | 30 | 30 | 30 | 30 | 30 | 30 | 30 |
| $p$ | — | 0.20 | 0.10 | 0.20 | 3.0 | 0.20 | 0.20 | 0.20 | 0.20 | 0.20 | 0.20 | 0.20 | 0.20 |
| $q$ | — | 0.10 | 0.10 | 0.10 | 0.10 | 0.10 | 0.10 | 0.10 | 0.10 | 0.10 | 0.10 | 0.10 | 0.10 |
| $N_a$ | ms | 125 | 60 | 125 | 63 | 125 | 200 | 40 | 125 | 125 | 125 | 125 | 125 |
| $N_T$ | ms | 15 | 15 | 15 | 40 | 15 | 300 | 10 | 75 | 75 | 75 | 75 | 75 |
| $(\alpha_T, \beta_T)$ | — | (9, 1) | (9, 1) | (9, 1) | (1, 1) | (9, 1) | (9, 1) | (9, 1) | (9, 1) | (9, 1) | (9, 1) | (9, 1) | (9, 1) |
| $(\alpha_c, \beta_c)$ | — | (9, 1) | (9, 1) | (9, 1) | (9, 1) | (9, 1) | (1, 1) | (1, 1) | (9, 1) | (9, 1) | (9, 1) | (9, 1) | (9, 1) |
| $\sigma_{\max}$ | px | 57.74 | 57.74 | 57.74 | 57.74 | 57.74 | 57.74 | 57.74 | 800 | 160 | 800 | 800 | 800 |
| $\sigma_{\min}$ | px | 57.74 | 57.74 | 57.74 | 57.74 | 57.74 | 57.74 | 57.74 | 40 | 8 | 40 | 40 | 40 |
| $r_0$ | px | 0 | 0 | 0 | 0 | 0 | 0 | 0 | 977 | 195 | 977 | 977 | 977 |
| $\Delta r$ | px | 0 | 0 | 0 | 0 | 0 | 0 | 0 | 61 | 12.2 | 61 | 61 | 61 |

### Experiments

We evaluated the ability of the proposed model to predict a wide range of motion phenomena, ranging from the basic to the as yet unexplained. For each of the 11 phenomena explored, we provide a brief description and detail the stimuli used in the tests.

1. *The missing-fundamental illusion* (Fig. S1). The missing-fundamental illusion occurs when removing the first harmonic from the Fourier series expansion of a square wave. This causes the pattern to be perceived as moving in the direction opposite to its physical displacement (Adelson and Bergen, 1985). Baseline control inputs were square-wave gratings displaced rightward at 6*°/*s in quarter-cycle steps. Missing-fundamental stimuli were generated by removing the fundamental harmonic from the baseline. Stimuli were evaluated at a spatial frequency of 1 cycle/°, spatial extent of 4°, and duration of 0.5 s.
2. *The reverse-Phi illusion* (Fig. S2). This is a visual phenomenon where inverting contrast polarity between successive displacement steps causes perceived motion in the direction opposite to the physical movement (Anstis,1970). Input sequences comprised random contrast patterns (Michelson contrast: 0.8) drifting at an effective velocity of 19.2*°/*s on a 0.5 s duration, with contrast polarity inverted at each frame transition for the reverse-Phi condition. Each condition was presented with 20 independent noise samples.
3. *Contrast saturation* (Fig. S3). Perceived motion strength scales with image contrast at low levels but saturates at higher contrast values (Heeger, 1992; Ohzawa et al., 1985; Albrecht and Hamilton, 1982). To evaluate model responses across contrast regimes, input sinusoidal gratings (spatial frequency: 1 cycle/°, velocity: 8*°/*s and duration: 0.5 s) were fed to the model across 20 logarithmically spaced contrast levels spanning [1%, 100%].
4. *Motion masking* (Fig. S4). Motion masking refers to the reduced detectability of a moving target signal when overlaid by a moving background masker sharing similar spatial frequencies (Anderson and Burr, 1985). Target stimuli were drifting sinusoidal gratings (Michelson contrast: 0.8, spatial frequency: 2.5 cycles/°, velocity: 10*°/*s) presented alongside a jittering mask grating (Michelson contrast: 0.8, temporal frequency: 10 Hz, randomized direction) across 13 mask spatial frequencies in the range [0.02, 12] cycles/°.
5. *Second-order motion* (Fig. S5). Second-order motion consists of drifting local contrast variations without corresponding shifts in mean luminance over space (Chubb and Sperling, 1988; Cavanagh and Mather, 1989). Stimuli were generated by modulating a static random-noise carrier texture with a drifting sinusoidal contrast envelope (spatial frequency: 1 cycle/°, velocity: 8*°/*s, duration: 0.5 s) moving rightward or leftward. One hundred independent carrier samples, each with a random envelope phase, were presented per direction.
6. *Direction of motion for gratings* (Fig. 1). The model was evaluated using drifting sinusoidal luminance gratings, each moving in the direction orthogonal to its bars (Adelson and Bergen, 1985). Stimuli (spatial frequency: 0.25 cycles/°, temporal frequency: 10 Hz, duration: 2 s) were presented at seven motion directions, spanning 0° to 90° in 15° increments.
7. *Direction of motion for Type I plaids* (Fig. 2). Type I plaids are 2D patterns formed by superimposing two sinusoidal gratings, configured such that vector average (VA) and intersection-of-constraints (IOC) yield identical motion direction (Adelson and Movshon, 1982; Ferrera and Wilson, 1990). Component gratings (spatial frequency: 1 cycle/°, equal speeds) were oriented at ±45° relative to the intended pattern trajectory. Pattern motion was evaluated across seven directions spanning 0° to 90° in 15° increments with a duration of 2 s.
8. *Direction of motion for Type II plaids* (Fig. 3). Type II plaids are 2D composite patterns where VA and IOC yield divergent perceived directions (Ferrera and Wilson, 1990), with human perception following the IOC under the present conditions of long duration and high contrast (Yo and Wilson, 1992). Stimuli were constructed from two sinusoidal component gratings (spatial frequency: 2 cycles/°) moving at a 2 : 1 speed ratio and a duration of 2 s. Seven orientation configurations were evaluated, yielding theoretical IOC pattern motion directions spanning −30° to +30° in 10° increments.
9. *Interaction among scales* (Fig. 4). Scale-dependent motion interactions can induce an illusory reversal in the perceived direction of a fine-scale component when it is added to a static coarse-scale pattern. This striking perceptual phenomenon typically occurs when the complex stimulus is presented at short durations. Stimuli were constructed by combining a static coarse-scale component (1 cycle/°) with a drifting fine-scale component (3 cycles/°, temporal frequency: 12 Hz, duration: 0.5 s). To evaluate the temporal dependence of the illusory motion reversal, stimuli were windowed by Gaussian temporal envelopes with widths from 10 ms to 200 ms. Each temporal duration was presented 25 times, with an independent random spatial phase for each of the two components.
10. *Breaking of time-reversal symmetry* (Fig. 5). Time-reversal asymmetry occurs when the magnitude of the visual system’s behavioral or motion response to a forward-playing sequence fails to match the opposite magnitude of its time-reversed counterpart. Following the paradigm of (Wu et al., 2025), the model was evaluated using two periodic spatial profiles presented in forward order and in exact frame-reversed temporal order (field of view: 3.52°): (1) a sawtooth edge pattern containing a single sharp luminance transition per cycle (spatial period: 1.17°, velocity: 9.38*°/*s, Michelson contrast: 1.0, duration: 1 s), and (2) an uneven square-wave grating characterized by asymmetric spatial duty cycles (spatial period: 2.34°, velocity: 1.17*°/*s, Michelson contrast: 1.0, duration: 1 s). For each of 40 random spatial phases, the same frame sequence was presented in forward order and reversed in time. Model responses were compared between the original frame sequences and their exact time-reversed counterparts to quantify non-linear deviations from temporal symmetry (Fig. 5). To quantify the breaking of time-reversal symmetry, we followed the criterion of (Wu et al., 2025) that a time-reversal symmetric response must satisfy *R*_fwd_ = − *R*_rev_, where *R*_fwd_ and −*R*_rev_ are the model responses to a stimulus played forward and in reverse, respectively. We define the antisymmetric and symmetric components of the response as

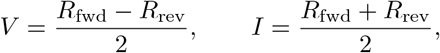

where *I* is the component that reflects the breaking of the symmetry (*I* = 0 for a perfectly symmetric response). The ratio *V/I* is the directional response relative to the symmetry-breaking component; thus, a smaller |*V/I* | means a stronger departure from time-reversal symmetry.
11. *Static motion illusions* (Fig. 6, Table 3). Static motion illusions are unmoving patterns that elicit illusory motion through asymmetric luminance profiles. Three illusions from (Rosario et al., 2026) were tested, and in all cases, the stimuli were presented with an abrupt temporal onset while remaining fully static in space: (1) *Rotating snakes*: an off-center pattern built from a repeating unit with an asymmetric sequence of luminance steps (black, dark gray, white, and light gray) (Kitaoka and Ashida, 2003; Murakami et al., 2006); (2) *Central drift* : a centered ring of elements whose luminance changes as a continuous asymmetric gradient from dark to light, the perceived motion following that gradient (Kitaoka and Ashida, 2004); (3) *Peripheral drift* : a repeating sawtooth luminance profile, a linear rise from black to white followed by an abrupt return to black, presented at the same off-center position as the rotating snakes (Faubert and Herbert, 1999; Kitaoka and Ashida, 2003). Two controls were also tested: a *symmetric control*, a stimulus with a symmetric luminance profile that serves as a control, since it lacks the asymmetric luminance gradient responsible for driving illusory motion, and a *real rotation* of that symmetric stimulus, rotating clockwise at 2.5*°/*s from the first frame, without onset. For the static motion illusions, we quantify how well the estimated flow matches the rotation perceived by human observers with the normalized correlation of (Rosario et al., 2026). Let *P* ∈ ℝ^*K*×2^ be the flow predicted by the model at *K* spatial locations, whose *i*-th row is the model output (**C***^H^,* **C***^V^*) at location *i*, and let *R* ∈ ℝ^*K*×2^ be the flow of a rigid rotation in the direction perceived by human observers, which is clockwise for the illusions considered here. Its magnitude grows with the distance *r_i_* to the center of the stimulus as (*r_i_/r*_out_)*^η^M*, where *r*_out_ is the radius of the stimulus, *M* the magnitude at its boundary, and *η* = 1. The alignment is

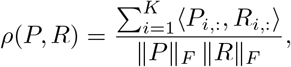

where ⟨·,·⟩ is the standard inner product in ℝ^2^ and ∥ · ∥*_F_* the Frobenius norm. The score lies in [−1, 1]: *ρ* = 1 indicates perfect agreement with the human percept, *ρ* = −1 rotation in the opposite direction, and *ρ*≈ 0 no rotation.

**Figure 1.**
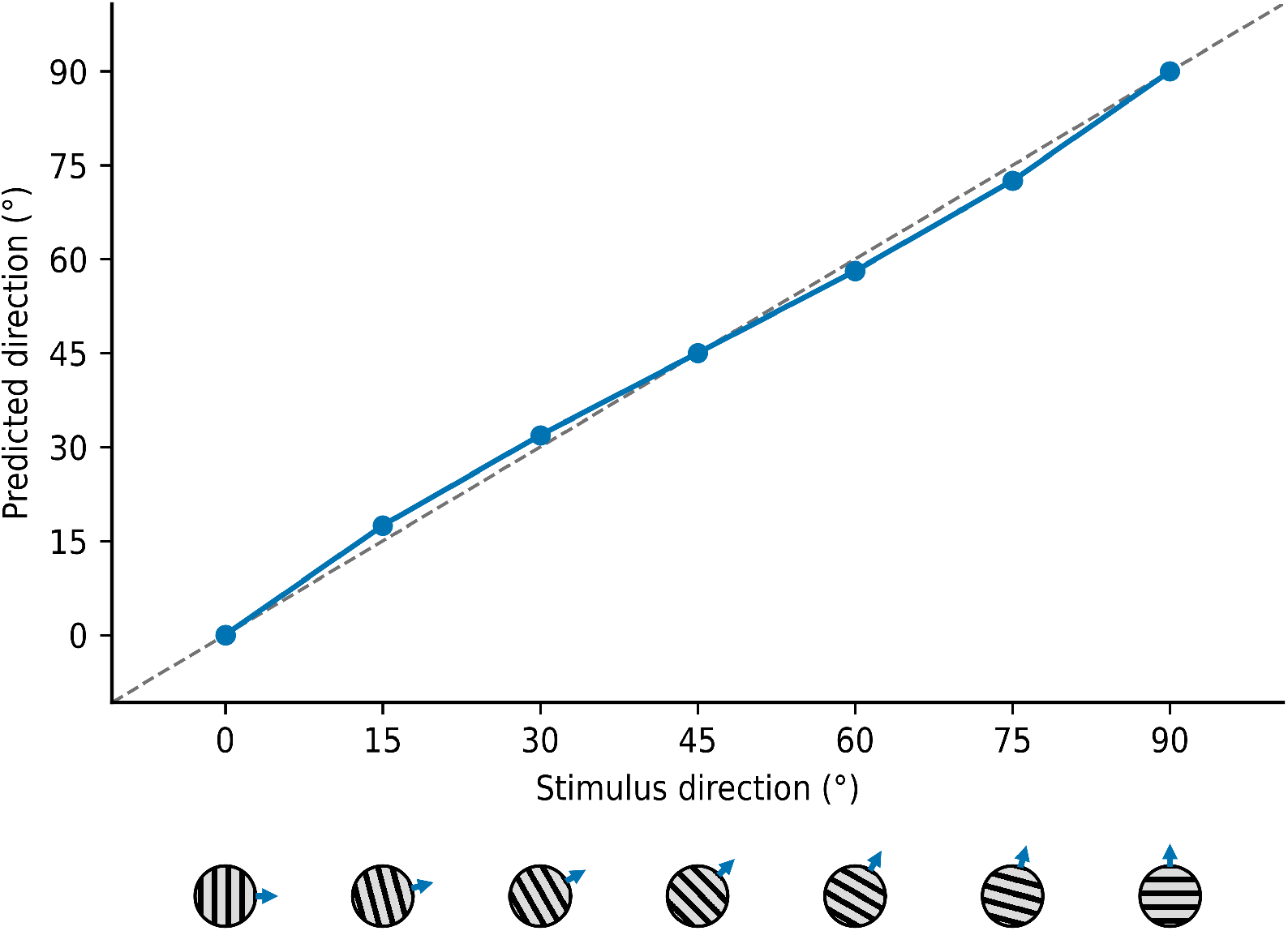
The model estimates the direction of moving gratings. Predicted direction as a function of the true direction of motion of a drifting sinusoidal grating, whose orientation and direction are shown by the icons below the horizontal axis. The dashed line marks perfect agreement between prediction and stimulus, which the model follows closely, with a mean absolute error of 1.2°.

**Figure 2.**
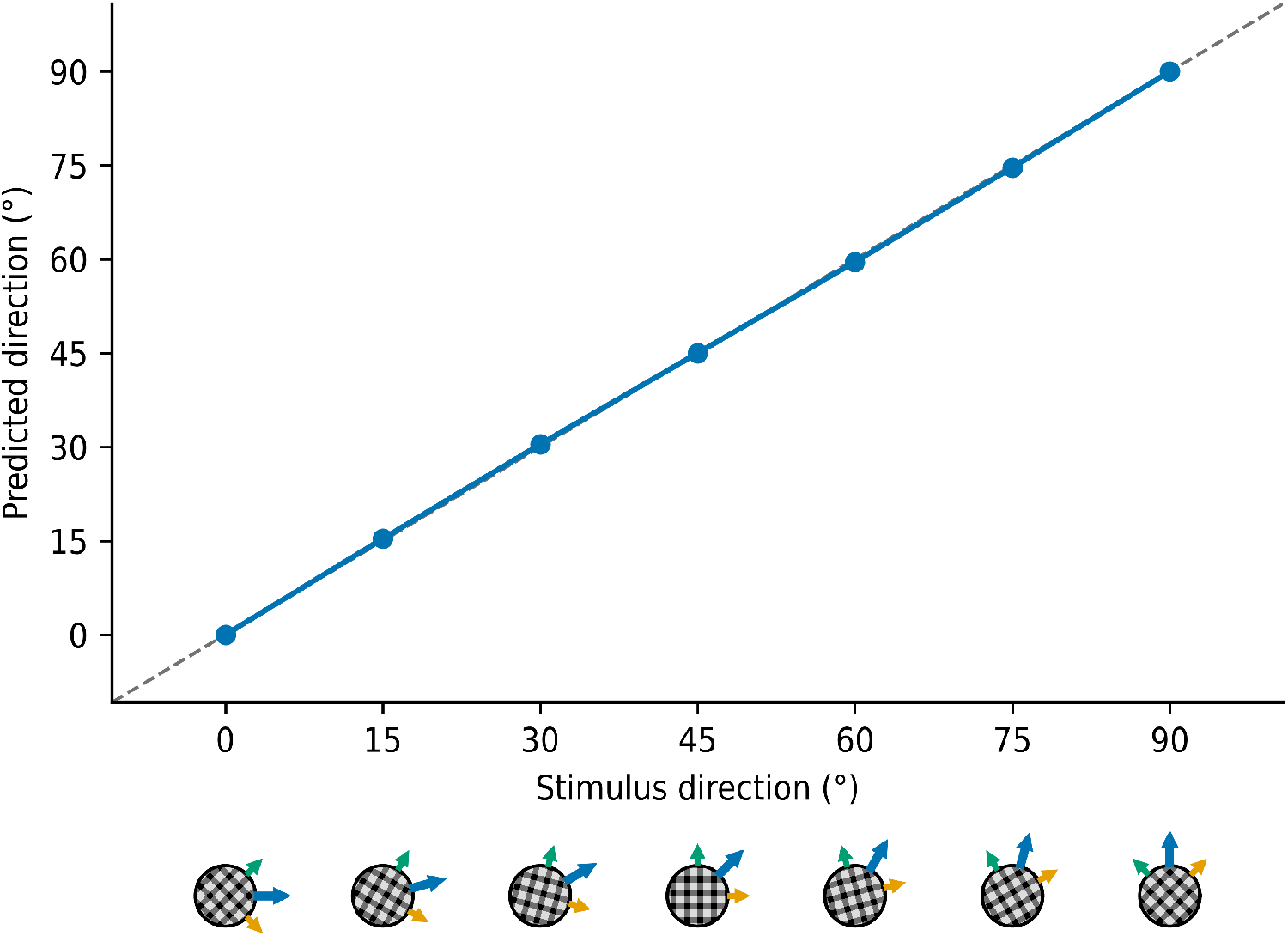
The model estimates the direction of moving Type I plaids. Predicted direction as a function of the pattern direction of a plaid formed by two sinusoidal gratings oriented at ±45° from that direction, a configuration for which the vector average and the intersection of constraints coincide. The icons show the plaid together with the directions of motion of its two components (green and orange arrows) and of the pattern as a whole (blue arrow); the dashed line marks perfect agreement, which the model reproduces closely, with a mean absolute error of 0.2°.

**Figure 3.**
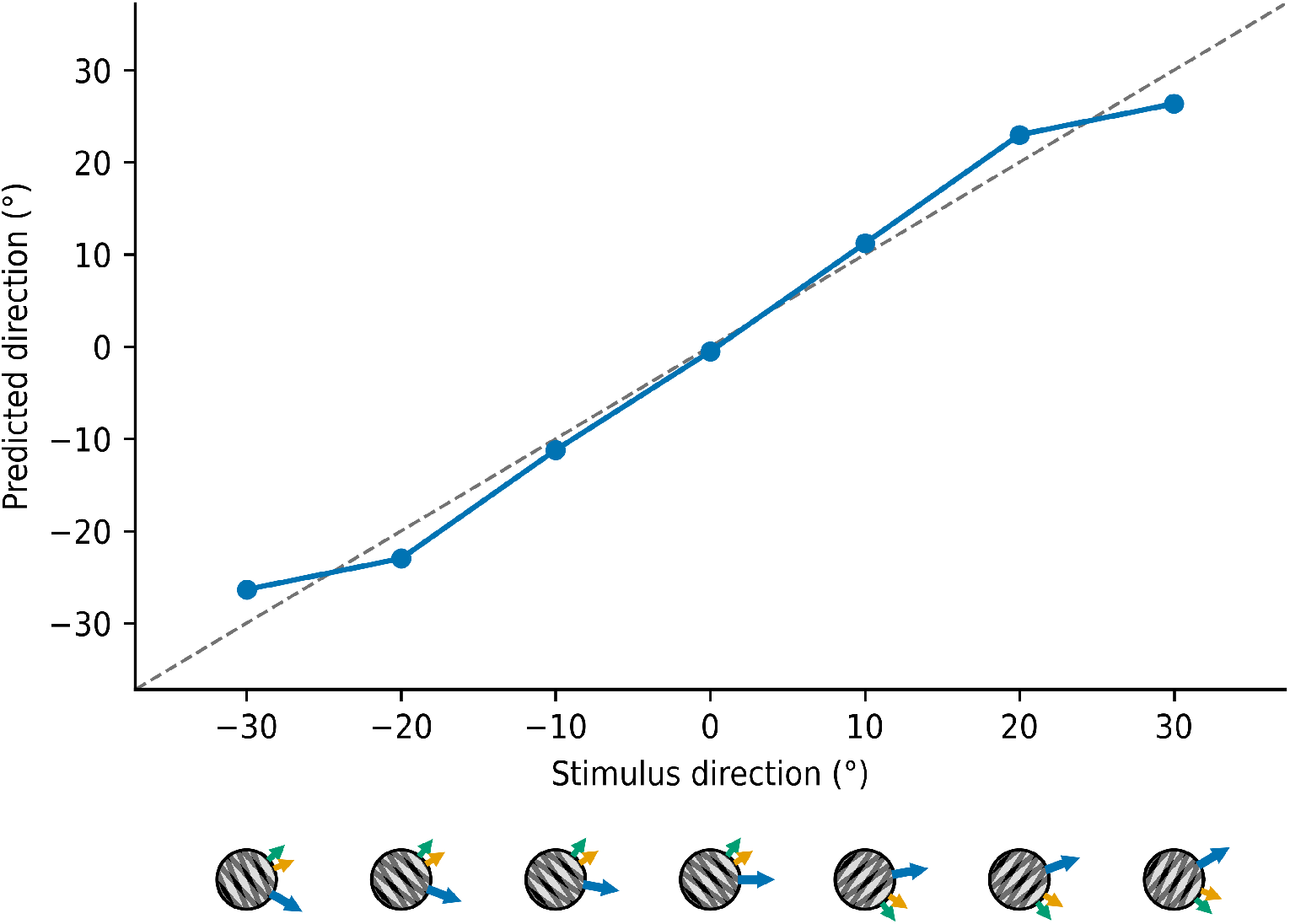
The model follows the intersection-of-constraints (IOC) direction for Type II plaids, for which the vector average and the IOC directions diverge. Predicted direction as a function of the IOC direction of the plaid; icons as in Fig. 2. The model tracks the IOC direction closely, with a mean absolute error of 2.3°.

**Figure 4.**
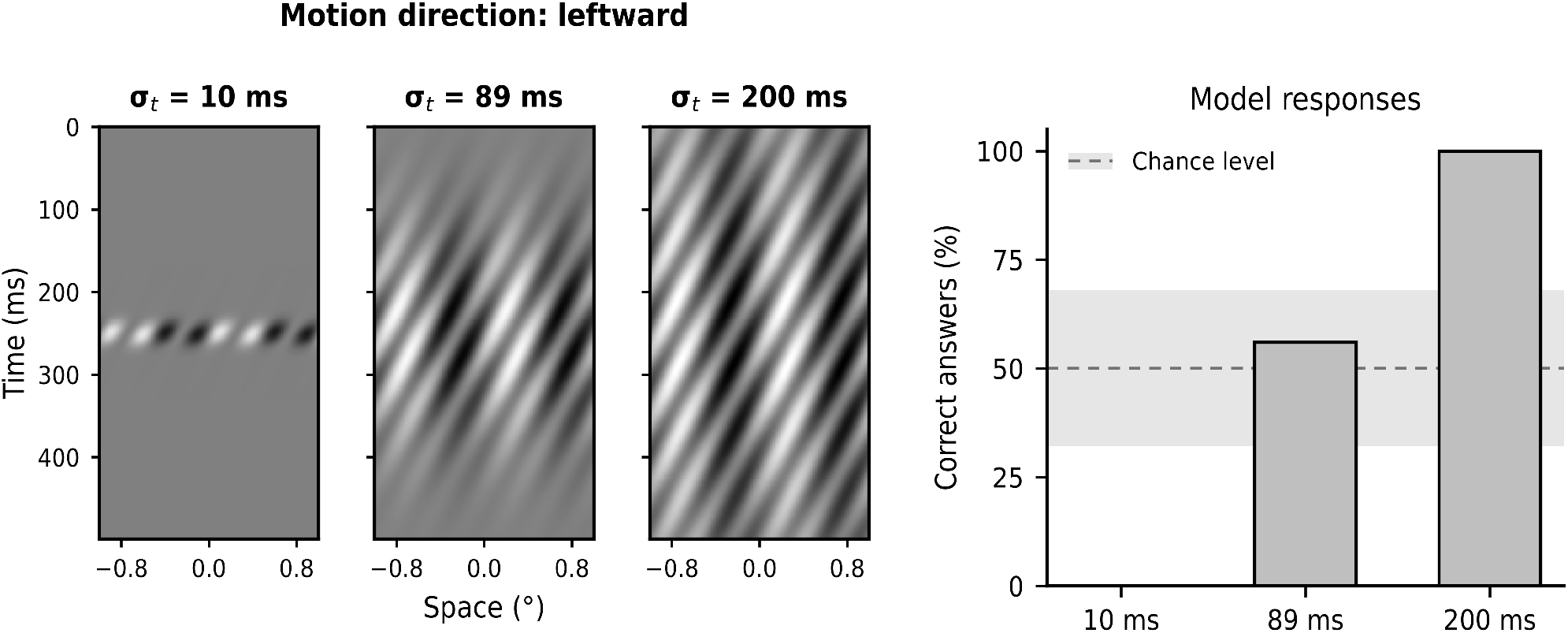
The model reproduces the inversion of perceived direction produced by the interaction among scales. The stimulus is the sum of a static coarse grating and a fine grating drifting leftward, windowed in time by a Gaussian temporal envelope whose duration is expressed as its standard deviation *σ_t_*. Left: space-time plots of the stimulus for *σ_t_* = 10, 67.5 and 100 ms. Right: percentage of presentations for which the model reports the physical, leftward direction, as a function of *σ_t_*, over 25 presentations with the spatial phase of each component drawn at random. The dashed line marks the performance expected by chance and the shaded band the limits, assuming binomial variability, within which performance cannot be distinguished from chance at the 5 per cent level. For the briefest presentations the model consistently reports the direction opposite to the physical one, and for the longest it reports the physical direction.

**Figure 5.**
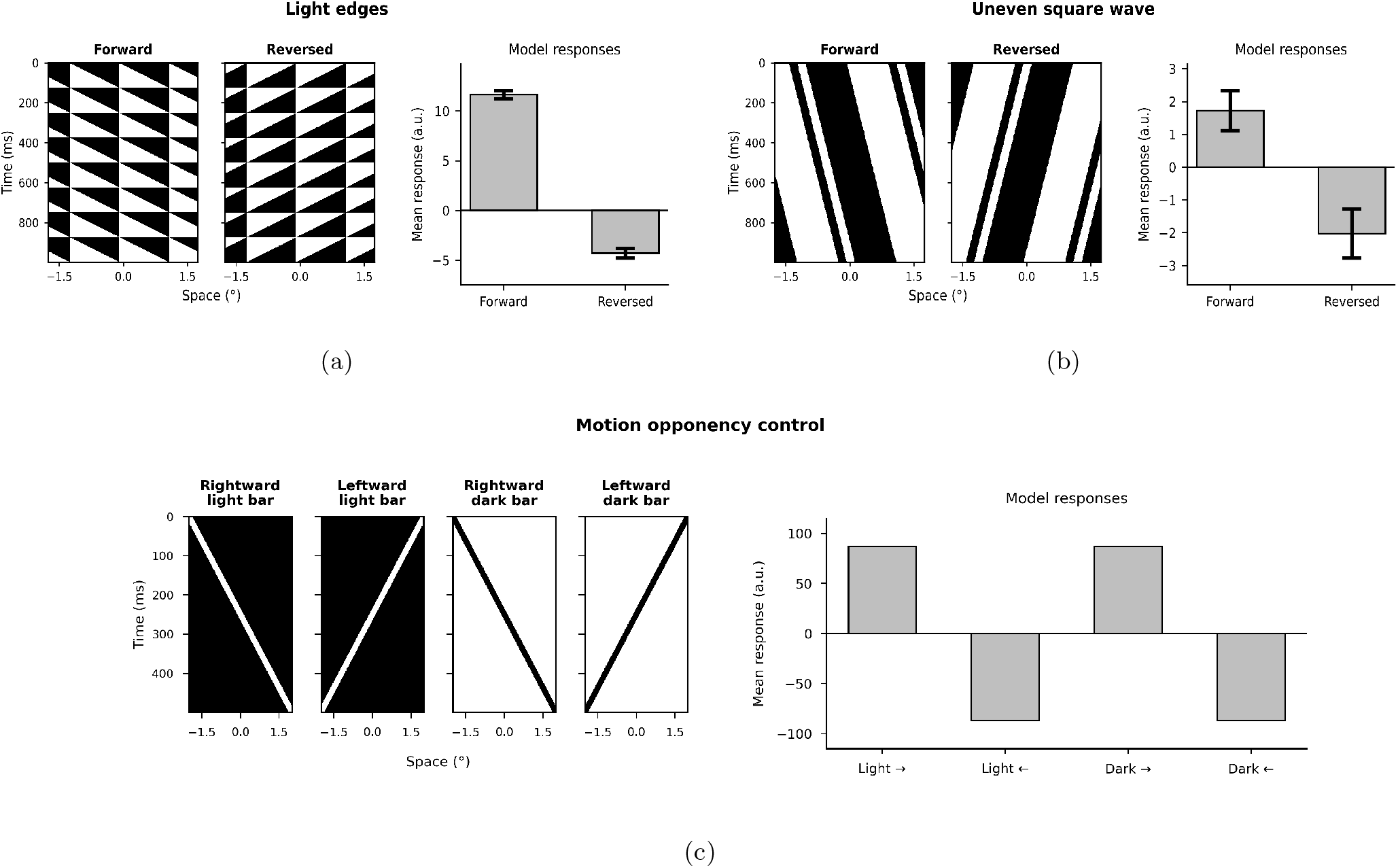
The model breaks time-reversal symmetry for the two stimuli designed to probe it, but not for the motion-opponency control. In each panel, the space-time plots show the stimulus played forward and in exact frame-reversed order, and the bar plot shows the mean model response to each; a time-reversal symmetric response would satisfy *R*_fwd_ = −*R*_rev_, so that the symmetric component *I* = (*R*_fwd_ + *R*_rev_)*/*2 vanishes and the ratio *V/I* diverges. (a) Light edges: the model responds with *R*_fwd_ = +11.62 and *R*_rev_ = −4.31 model units, giving *V/I* = +2.18. (b) Uneven square wave: *R*_fwd_ = +1.72 and *R*_rev_ = −2.03, giving *V/I* = −12.27. (c) Motion opponency, used here as a control: a light or a dark bar moving rightward or leftward elicits responses of equal magnitude and opposite sign (±87.18 for both polarities), so that *I* is negligible, as expected for a genuinely symmetric response. In (a) and (b), responses are averaged across 20 presentations with varying spatial phase of the stimuli.

### Comparison with DNNs for motion estimation

We evaluated the five motion-estimation DNN models listed in Table 2. All of them estimate dense optical flow, and they differ in how much of their design is motivated by biology. Three of them, PWC-Net, LiteFlowNet2, and RAFT, estimate flow from a pair of frames by matching learned features rather than raw intensities, on feature maps of lower resolution than the input. PWC-Net (Sun et al., 2018) follows a coarse-to-fine scheme over a feature pyramid: at each level, it warps the features of the second frame by the current estimate, builds a partial cost volume against those of the first frame, and a CNN predicts the refined flow, which is passed to the next level. LiteFlowNet2 (Hui et al., 2021) uses the same pyramidal scheme with a lightweight design, combining cascaded flow inference through feature matching and sub-pixel refinement with a learned flow regulariser at each level. RAFT (Teed and Deng, 2020) departs from coarse-to-fine refinement: it extracts per-pixel features from the two frames, builds a correlation volume containing all pairwise feature similarities, and updates a single flow field, kept at a fixed resolution, through a recurrent operator that iteratively looks up values in that volume. The other two DNNs are biologically inspired, in that their components are motivated by stages of the visual system, and they take a sequence of frames rather than a pair. ME-Attention (Sun et al., 2023) has two stages: a trainable motion-energy front end, which extracts local motion signals with spatio-temporal filters as V1 cells do, and a self-attention network that integrates those signals over space and time. Dual (Sun et al., 2025) extends that architecture into a two-pathway system, one pathway for luminance-defined motion and another for contrast-defined motion, so that the model accounts for both first- and second-order motion.

**Table 2.** The five motion-estimation models evaluated. Parameter counts are those of the released weights used here; input size and frames per inference are each model’s native configuration.

| Model | Parameters | Input size (px) | Frames per inference |
| --- | --- | --- | --- |
| PWC-Net (Sun et al., 2018) | 9.4 M | 1024 × 436 | 2 |
| LiteFlowNet2 (Hui et al., 2021) | 6.4 M | 1024 × 436 | 2 |
| RAFT (Teed and Deng, 2020) | 5.3 M | 960 × 520 | 2 |
| ME-Attention (Sun et al., 2023) | 14.7 M | 768 × 768 | 11 |
| Dual (Sun et al., 2025) | 25.6 M | 768 × 768 | 15 |

**Figure 6.**
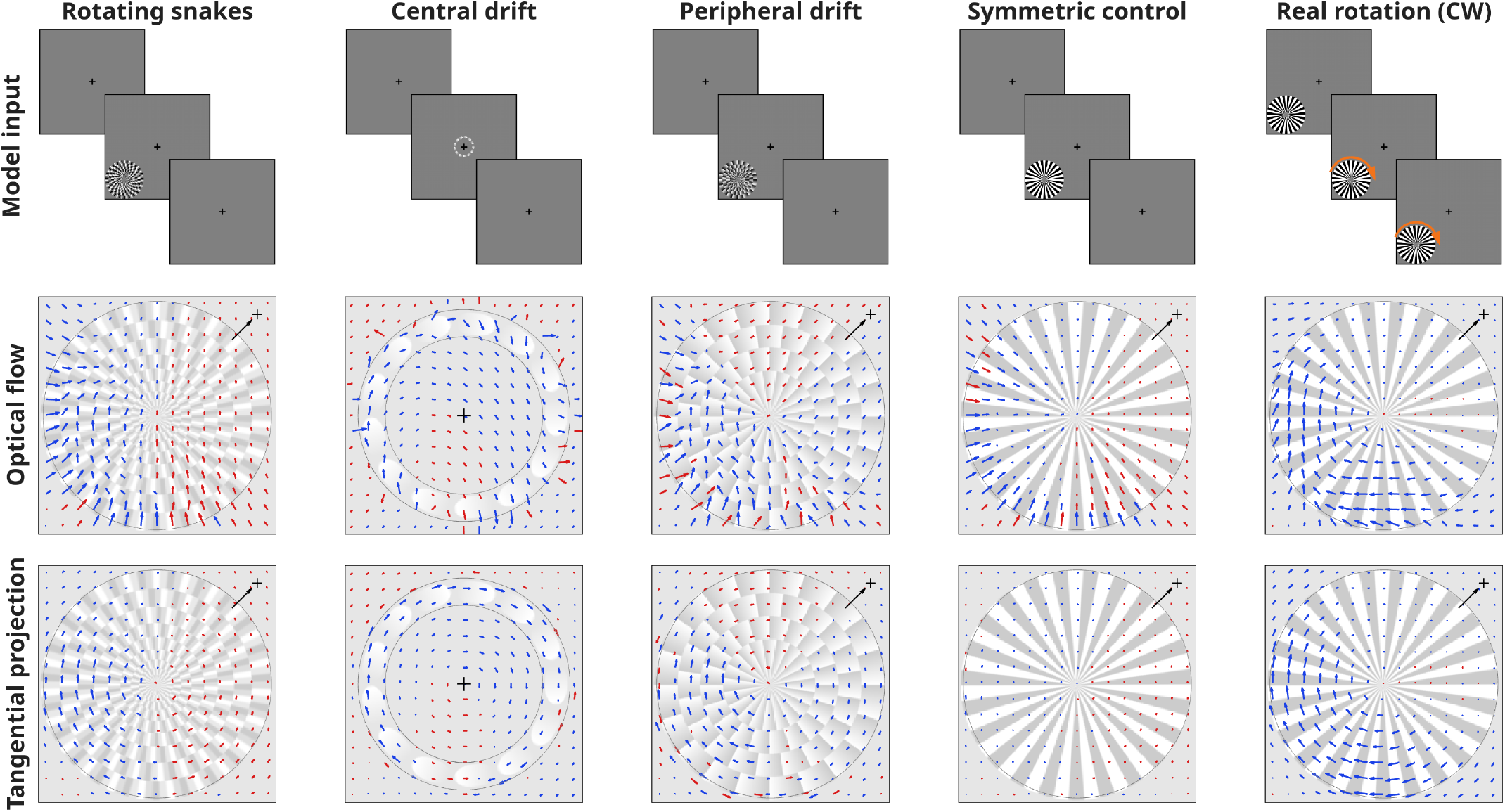
The model produces an illusory rotation in the direction perceived by human observers for all three static motion illusions tested. Columns, left to right: rotating snakes, central drift and peripheral drift, followed by two controls: a stimulus with a symmetric luminance profile that serves as a control, since it lacks the asymmetric gradient that drives illusory motion, and that same symmetric stimulus pattern physically rotating clockwise. Top row: three frames of the model input; the stimulus appears abruptly after a blank interval and disappears again, except for the real rotation, which is present from the first frame and whose direction is indicated by the orange arrow. The cross marks the center of the canvas, which is the fixation point. Middle row: optical flow estimated by the model. Bottom row: the tangential component of that flow with respect to the center of the stimulus, arrows are blue where this component is clockwise and red where it is counterclockwise, and their length is normalized within each column. The black arrow in the corner of each map points toward the fixation point, and the gray circle delimits the region of the stimulus over which the alignment *ρ* is computed.

To ensure a fair comparison, all models were evaluated using the official public implementations and pretrained weights released by their respective authors, with their native inference settings: the input length listed in Table 2, that is, the number of frames each model takes per inference, inputs at each model’s required resolution and intensity range, and grayscale frames replicated across the three color channels. No fine-tuning was performed.

The networks were run on the same experiments as our model, with the same stimuli, conditions, and number of repetitions (Experiments 1–11), but in order to ensure a fair comparison the sampling of the stimuli (pixels per degree and framerate) was adapted so as to guarantee that average speed and spatial frequency fell within the value ranges at which the networks have been shown to perform successfully using naturalistic video inputs. Two constraints were applied. First, the displacement between consecutive frames was kept between 10 and 40 px: below that range the motion is lost when the input is reduced in resolution inside the network, and above it the correspondence falls outside the search range these architectures provide. Second, for periodic stimuli the displacement was kept at or below a quarter of the spatial period, well below the half period at which the direction of motion becomes ambiguous. Conditions that are compared with each other were rendered with the same pixels per degree and the same frame rate, so that the displacement between frames is identical across them and any difference in the response can only be attributed to the variable under study, such as contrast or mask frequency, and not to a change in sampling.

Whole stimulus sequences were processed: the two-frame models were run on every consecutive pair of frames and the multi-frame models on sliding windows of their input length, so that each model returns a dense optical flow field per pair or per window. These flow fields were then reduced to a single motion vector per stimulus with the third stage of our model (Eq. 3), in the configuration used for that same experiment with our model: spatial pooling with *c_s_* over a region covering the same visual angle, and temporal pooling with *c_t_* using the same parameters. Applying the same readout used for our model to the networks makes the two sets of estimates directly comparable. We report the direction 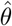 of the resulting vector and its magnitude. Across repetitions we report the circular mean of 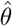, its circular standard deviation, and the mean absolute error with respect to the direction of the physical motion, computed over the individual repetitions.

## Results

Given that the highly nonlinear nature of our model impedes its compact characterization as a cascade of linear filters and pointwise nonlinearities, as is commonly done in standard approaches, we have decided to call our model INME, for intrinsically nonlinear motion estimation. We begin the experiments by verifying that INME is able to reproduce the same motion phenomena reported for the 1D motion sensor model of (Luna et al., 2025) (of which the second stage of our model is an extension) and which have been observed in insects and vertebrates: first and second-order motion, the missing-fundamental and reverse-Phi illusions, contrast saturation and motion masking (Experiments 1–5, and the opponency control of Experiment 10). This is indeed the case, as shown in Fig. 5c and in the Supplementary Material (Figs. S1–S5). Of note, existing first-order motion models are not well suited to detect second-order motion without additional processing; in contrast, the model proposed here and the 1D motion sensor in (Luna et al., 2025) detect both first- and second-order motion while preserving the exact same form, without the need to perform structural changes or parameter modifications when switching between motion types.

Figs. 1, 2 and 3 show that the model is able to estimate the direction of moving gratings and the direction of moving plaids of both Type I and II (Experiments 6–8); for Type II plaids, Fig. 3 shows that the direction predicted by the model closely matches the intersection-of-constraints direction which agrees with data from neurophysiology and psychophysics. Again, these are basic phenomena observed in a wide variety of animals and visual system stages with which our model would be compatible.

For Experiment 9, *interaction among scales*, we see in Fig. 4 that the proposed model predicts the inversion of perceived direction at short stimulus durations of a complex stimulus composed of a static, low spatial frequency grating added to a moving high spatial frequency grating. This is a puzzling phenomenon reported only in humans (Derrington and Henning, 1987) and explained in the literature by a single model (Serrano-Pedraza et al., 2007), that accounts for several aspects of scale interaction (Serrano-Pedraza and Derrington, 2010; Luna and Serrano-Pedraza, 2018; Bachtoula and Serrano-Pedraza, 2025) but which cannot account for the other motion phenomena examined in the present study, as it was specifically developed to explain the interaction effect and not intended as a general model of motion perception.

For Experiment 10, the breaking of time-reversal symmetry, Fig. 5 shows that the proposed model predicts results that have been recently reported in the fly and are not explained by any established vision motion estimation model (Wu et al., 2025).

Finally, for the three static motion illusions that we tested (Experiment 11), the model predicts a rotation in the direction perceived by human observers (Fig. 6): the alignment *ρ* between the optical flow predicted by the model and the clockwise rotation perceived by observers is positive for the rotating snakes, the central drift and the peripheral drift, close to zero for the symmetric control, and highest for the physically rotating pattern (Table 3). Neural models for these types of illusions, reported for insects and vertebrates, are usually developed just to explain peripheral drift and do not consider other types of motion phenomena (Agrochao et al., 2020; Bach and Atala-Gérard, 2020).

**Table 3.** Alignment *ρ* between the optical flow estimated by the model and the clockwise rotation perceived by human observers, computed over the whole stimulus. Positive values indicate agreement with the human percept, and the closer *ρ* is to 1, the stronger that agreement.

| Stimulus | $\rho$ |
| --- | --- |
| Rotating snakes | +0.147 |
| Central drift | +0.132 |
| Peripheral drift | +0.174 |
| Symmetric control | +0.002 |
| Real rotation (CW) | +0.632 |

In short, these results show that our model is indeed able to predict all the motion phenomena considered. For comparison, we also tested five state-of-the-art DNNs for motion estimation, some of them bio-inspired and all of them having millions of parameters. We found that, when presented with the same synthetic stimuli used to test our model, these DNNs are only capable of predicting a few of the contemplated phenomena, and none of them is able to predict interaction among scales, the breaking of time reversal symmetry nor the three motion illusions analyzed. Table 4 summarizes which phenomena each network reproduces; the individual measurements behind it are given in the Supplementary Material (Tables S1–S11).

**Table 4.** Results of the five DNN optical flow models across the eleven psychophysical experiments considered in this work. A checkmark indicates that the model reproduces the phenomenon, a cross that it fails to do so, and a tilde a partial or ambiguous result.

| Experiment | RAFT | PWC-Net | LFN2 | ME-Att. | Dual |
| --- | --- | --- | --- | --- | --- |
| The missing-fundamental illusion | × | × | × | ✓ | × |
| The reverse-Phi illusion | × | × | × | × | × |
| Contrast saturation | × | × | × | × | × |
| Motion masking | × | × | × | × | × |
| Second-order motion | × | × | × | ✓ | ✓ |
| Direction of moving gratings | × | × | × | × | × |
| Direction of moving plaids of Type I | × | × | × | × | × |
| Direction of moving plaids of Type II | × | × | × | × | ✓ |
| Interaction among scales | × | × | × | × | × |
| Breaking of time reversal symmetry | × | × | × | × | × |
| Static motion illusions<sup>a</sup> | × | × | × | × | × |

<sup>a</sup> Taken from (Rosario et al., 2026).

## Discussion

Here we have introduced a neural circuit model that only uses cells selective to cardinal directions and is able to predict a wide range of visual motion data, making it a good candidate for a canonical neural circuit for motion direction estimation. As a possible mechanistic interpretation of our model, we propose that the linear spatial terms would correspond to passive dendrites, and the nonlinear function *f* (·) would account for the response of active dendrites, with the subtracting element in its argument representing a back-propagating action potential (bAP). The importance of dendritic nonlinearities for DS cells responsive to cardinal directions has been remarked upon by a number of recent works (Currier et al., 2023; DePiero et al., 2024; Kühn et al., 2025), and models that combine active dendrites with bAPs have shown remarkable capabilities in predicting very challenging experimental data from visual psychophysics (Bertalmío et al., 2024) and neurophysiology (Rentzeperis et al., 2025), and for creating DNNs that are more robust and data-efficient (Mohedano et al., 2026). Our reasoning is that the power of our model lies in its *joint* emulation of dendritic nonlinearities and bAPs, something that to the best of our knowledge has been absent until now from motion direction estimation models and that provides our approach with a greater flexibility and expressive power, allowing for dynamic changes in the neuron’s properties in response to complex stimuli.

This analysis, where the highly nonlinear nature of our model is what endows it with higher expressivity than models consisting of cascades of linear filters and pointwise nonlinearities, is supported by the fact that the state-of-the-art DNNs for motion estimation, each of them having millions of parameters, are not able to reproduce most of the visual phenomena that our model can predict while using just a small number of hand-crafted elements described by a few parameters in total. Some of the failures of these DNNs are due to the fact that they have been trained on videos emulating natural visual stimuli, where richly textured image regions are the norm but synthetic stimuli like sinusoidal gratings are scarce or absent. For example, all five DNNs perform poorly with “clean” sinusoidal gratings, but their performance is excellent when textured noise is added to the input (Fig. S6); this dependence on texture has already been reported for DNNs trained on natural photographs, which are known to rely on texture rather than shape and to fail when that cue is altered or removed (Geirhos et al., 2019). On the other hand, the proposed model yields an accurate estimate of the motion direction of the grating, regardless of the absence or presence of a superimposed texture. In any case, the results we report for motion estimation DNNs throughout this work (Table 4) show that these DNNs cannot be considered vision models, as they fail to replicate very basic properties of motion vision.

The purpose of the experiments has been to show that our proposed model can qualitatively replicate a wide variety of experimental data, and with that goal in mind we have not strived for accurate quantitative matching. However, we did observe that for several basic phenomena a single set of model parameter values is sufficient (this was the case for the missing-fundamental illusion, contrast saturation, motion masking and reverse-Phi), while more challenging experiments have required different parameter values, which suggests that different phenomena might involve different stages or areas of the visual system, and/or different types of cells.

Related to this, we want to stress a fundamental point: although we consider our model to be a good candidate for a canonical neural circuit, we do not claim nor believe that it encompasses *all* aspects involved in motion direction estimation, i.e. that for motion computation the visual system *only* uses a circuit like the one proposed here. The experimental results we obtain simply suggest that our model is enough to explain, qualitatively, the occurrence of varied motion phenomena on the retina and different brain regions, but other neural computations can also be influencing the outcome, or even producing a similar outcome through a different mechanism. This is consistent with the fact that, in mammals, there are two parallel motion pathways from the retina to the cortex, one carrying signals from cardinally-tuned DS cells in the retina, the other generating direction selectivity in V1 from non-direction-selective inputs from LGN (He et al., 2026), but it remains to be established whether or not direction selectivity computed *de novo* is combined, at some stage, with direction selectivity coming from the retina (Lien and Scanziani, 2018).

Our results also show that there is room for improvement for the model: for instance, in the case of interaction among scales, the qualitative response of the model is completely flat when there is no inversion, which does not accurately represent the experimental data. We are currently investigating a number of avenues in order to extend our model and strengthen its predictive power: (1) analyzing the mathematical properties of the model and its connections with efficient representation and predictive coding approaches (Clark and Fitzgerald, 2024); (2) adding terms corresponding to other dendritic nonlinearities; (3) considering also DS cells in arbitrary, not just cardinal, directions; (4) studying the role of image and noise statistics in the breaking of time-reversal symmetry, as observed in (Meding et al., 2019) and (Wu et al., 2025); (5) testing the model against a more extensive set of visual motion phenomena, including other challenging visual illusions like the Pinna-Brelstaff, Expanding hole and Enigma illusions (Pinna and Brelstaff, 2000; Laeng et al., 2022; Zeki et al., 1993), as well as fundamental motion phenomena such as induced motion (Duncker, 1929), motion surround suppression (Tadin et al., 2003), motion capture (Ramachandran and Cavanagh, 1987), the motion aftereffect (Mather et al., 2008), and speed perception (Stone and Thompson, 1992).

We also plan on moving from the hand-crafted elements of the current instance of our model (spatial and temporal filters, linear and nonlinear functions, scalar weights) to a data-driven implementation where the model elements are described as sets of weights that are learned so as to fit experimental data, in a machine learning approach. This methodology would provide higher predictive accuracy, facilitate the generation of functional hypotheses and allow us to synthesize stimuli with which neurons can be characterized (Walker et al., 2019). It also might suggest bio-inspired strategies to improve the performance and the properties of artificial neural networks: we are optimistic in this regard, given how our model is already outperforming DNNs trained for motion estimation.

## Code availability

The source code with the implementation of the model and the generation of all figures is available on GitHub: https://github.com/vmg-io-csic/INME

## Acknowledgements

This work has been supported by: Programa Fundamentos (ref. VIS4NN) of Fundación BBVA 2022 (Spain); Programa Tecnologías (ref. TEC 2024/COM-322) of Comunidad de Madrid (Spain); Proyectos de Generación de Conocimiento 2024 (ref. PID2024-161841NB-I00) of Ministerio de Ciencia, Innovación y Universidades (Spain); Programa Momentum (ref. MMT24-IO-02) of Consejo Superior de Investigaciones Científicas (Spain), funded by European Commission– NextGenerationEU through the Spanish Recovery, Transformation and Resilience Plan; and grant PID2024-155243NB-I00 (P.I. Ignacio Serrano-Pedraza) funded by MICIU/AEI/10.13039/501100011033 and by FEDER, UE.

## Author contributions

Conceptualization and methodology: Marcelo Bertalmío; numerical implementation: María Berraquero; experiments: María Berraquero and Ignacio Serrano-Pedraza; analysis of results and writing: all authors.

