## Supplementary Material: Figures S1-S6 and Tables S1-S10 for "A canonical neural circuit for motion direction estimation"

### Performance of state-of-the-art DNNs

Tables S1–S10 report the responses of the five DNNs (RAFT, PWC-Net, LFN2, ME-Attention, and Dual) to the stimuli described in the Methods section. The same quantities appear in all of them: the Direction column is the direction of the physical motion, $\hat{\theta}$ is the direction the network reports in degrees, and mag the magnitude of the vector it reports in pixels per frame. Where a condition was presented more than once, $\hat{\theta}$ is the circular mean across presentations and SD the circular standard deviation. The last row of each table gives the mean absolute error (MAE) of $\hat{\theta}$ with respect to the physical direction, taken over the individual presentations, and its standard deviation.

| Direction | RAFT | | PWC-Net | | LFN2 | | ME-Att | | Dual | |
|---|---|---|---|---|---|---|---|---|---|---|
| (°) | $\hat{\theta}$ (°) | mag | $\hat{\theta}$ (°) | mag | $\hat{\theta}$ (°) | mag | $\hat{\theta}$ (°) | mag | $\hat{\theta}$ (°) | mag |
| 0 | 1.8 | 16.55 | −58.5 | 29.50 | −51.7 | 26.03 | 19.4 | 12.01 | 14.3 | 19.52 |
| 15 | 5.7 | 16.42 | −68.6 | 138.77 | −29.9 | 23.28 | 10.4 | 6.22 | −19.8 | 21.99 |
| 30 | 84.7 | 28.07 | −54.8 | 167.40 | −11.6 | 21.09 | 41.7 | 6.63 | 20.3 | 21.28 |
| 45 | −11.3 | 27.42 | −40.7 | 210.79 | −36.5 | 106.69 | 59.6 | 7.31 | 73.4 | 20.08 |
| 60 | 102.4 | 22.57 | −33.6 | 92.11 | −29.8 | 136.34 | 46.3 | 3.58 | 71.0 | 21.68 |
| 75 | 66.6 | 17.52 | −25.9 | 90.39 | −10.0 | 102.43 | 72.1 | 16.59 | 74.9 | 15.31 |
| 90 | 107.8 | 16.44 | 6.1 | 122.16 | 171.6 | 104.93 | 93.7 | 8.44 | 80.3 | 16.23 |
| MAE ± SD | 27.3 ± 21.5 | | 84.4 ± 12.1 | | 68.0 ± 19.4 | | 10.1 ± 5.9 | | 15.4 ± 11.1 | |

Table S1: Moving gratings.

| IOC direction | RAFT | | PWC-Net | | LFN2 | | ME-Att | | Dual | |
|---|---|---|---|---|---|---|---|---|---|---|
| (°) | $\hat{\theta}$ (°) | mag | $\hat{\theta}$ (°) | mag | $\hat{\theta}$ (°) | mag | $\hat{\theta}$ (°) | mag | $\hat{\theta}$ (°) | mag |
| 0 | −24.4 | 199.07 | −120.2 | 32.24 | −0.3 | 12.11 | −29.4 | 6.96 | −2.0 | 10.99 |
| 15 | 132.0 | 26.98 | −95.7 | 77.66 | 14.1 | 10.63 | −14.5 | 4.48 | 5.8 | 11.72 |
| 30 | 30.1 | 12.12 | −65.1 | 44.65 | −91.0 | 35.18 | 47.2 | 15.07 | 14.8 | 9.48 |
| 45 | 44.8 | 12.02 | −84.2 | 43.73 | 166.7 | 34.99 | 29.1 | 11.35 | 14.4 | 36.71 |
| 60 | −61.0 | 35.20 | −81.8 | 74.62 | −178.9 | 35.09 | 67.6 | 12.15 | 55.5 | 14.75 |
| 75 | −99.7 | 44.35 | −104.3 | 47.96 | −164.2 | 33.27 | 56.3 | 9.29 | 80.8 | 12.56 |
| 90 | −26.9 | 107.21 | −96.9 | 59.03 | 89.6 | 12.03 | −145.9 | 30.79 | 98.4 | 8.85 |
| MAE ± SD | 79.2 ± 64.6 | | 135.6 ± 29.0 | | 69.5 ± 59.7 | | 34.6 ± 37.2 | | 10.8 ± 9.0 | |

Table S2: Type I plaids. The direction of the pattern as a whole is determined by the intersection of the constraints (IOC).

| IOC direction | RAFT | | PWC-Net | | LFN2 | | ME-Att | | Dual | |
|---|---|---|---|---|---|---|---|---|---|---|
| (°) | $\hat{\theta}$ (°) | mag | $\hat{\theta}$ (°) | mag | $\hat{\theta}$ (°) | mag | $\hat{\theta}$ (°) | mag | $\hat{\theta}$ (°) | mag |
| −30 | −29.8 | 14.01 | −60.5 | 20.79 | −29.9 | 13.59 | −92.2 | 3.39 | −25.7 | 15.96 |
| −20 | −177.6 | 54.48 | −23.5 | 14.36 | −55.5 | 26.90 | −90.4 | 3.41 | −14.6 | 14.44 |
| −10 | −10.6 | 13.44 | −54.3 | 42.15 | 106.0 | 44.83 | −62.6 | 6.85 | −2.5 | 15.23 |
| +0 | −0.8 | 13.63 | −21.8 | 18.40 | 34.3 | 135.94 | −62.2 | 1.22 | 3.9 | 13.61 |
| +10 | 124.1 | 261.02 | −59.6 | 48.68 | 44.4 | 135.17 | −44.2 | 1.51 | 9.2 | 12.81 |
| +20 | −40.8 | 545.65 | −50.7 | 51.86 | 38.2 | 136.05 | −45.1 | 1.51 | 27.3 | 13.00 |
| +30 | −30.2 | 334.88 | 32.8 | 13.89 | 28.0 | 137.44 | −30.1 | 2.24 | 25.4 | 13.93 |
| MAE ± SD | 56.3 ± 57.3 | | 34.7 ± 26.1 | | 34.4 ± 36.2 | | 61.0 ± 5.7 | | 4.8 ± 2.1 | |

Table S3: Type II plaids.

| Cond. | $\sigma_t$ (ms) | Direction | RAFT | | PWC-Net | | LFN2 | | ME-Att | | Dual | |
|---|---|---|---|---|---|---|---|---|---|---|---|---|
| | | (°) | $\hat{\theta}\pm$ SD (°) | mag | $\hat{\theta}\pm$ SD (°) | mag | $\hat{\theta}\pm$ SD (°) | mag | $\hat{\theta}\pm$ SD (°) | mag | $\hat{\theta}\pm$ SD (°) | mag |
| Moving 3 cpd | 10 | 180 | −23.8 ± 60.6 | 2.22 | −82.8 ± 0.0 | 182.97 | −56.3 ± 0.5 | 15.83 | 112.7 ± 57.5 | 0.32 | 52.6 ± 43.5 | 3.86 |
| + static 1 cpd | 25 | 180 | −45.6 ± 41.7 | 6.95 | −84.0 ± 0.1 | 200.12 | −53.4 ± 1.1 | 25.29 | −0.4 ± 0.1 | 1.62 | 109.0 ± 98.8 | 8.21 |
| | 37.5 | 0 | 39.6 ± 82.7 | 18.28 | −85.3 ± 0.2 | 222.67 | −59.3 ± 2.9 | 36.43 | −0.7 ± 0.1 | 2.99 | −71.3 ± 43.2 | 9.12 |
| | 100 | 0 | −74.4 ± 28.3 | 57.92 | −82.6 ± 1.0 | 239.52 | −13.7 ± 3.5 | 12.24 | −3.6 ± 0.2 | 4.08 | −4.7 ± 6.5 | 22.72 |
| MAE ± SD | | | 81.4 ± 31.6 | | 87.3 ± 5.8 | | 62.5 ± 39.5 | | 19.0 ± 40.5 | | 73.7 ± 63.9 | |

Table S4: Interaction among scales. $\sigma_t$ is the width of the temporal envelope, and Cond. identifies which component moves. The values were averaged across 25 presentations with varying spatial phases of each component.

| Contrast | Direction | RAFT | | PWC-Net | | LFN2 | | ME-Att | | Dual | |
|---|---|---|---|---|---|---|---|---|---|---|---|
| | (°) | $\hat{\theta}$ (°) | mag | $\hat{\theta}$ (°) | mag | $\hat{\theta}$ (°) | mag | $\hat{\theta}$ (°) | mag | $\hat{\theta}$ (°) | mag |
| 0.0100 | 0 | 34.4 | 12.43 | −69.5 | 38.27 | −20.1 | 99.69 | −9.3 | 1.45 | 1.2 | 11.01 |
| 0.0127 | 0 | −0.1 | 10.06 | −70.7 | 38.48 | −19.9 | 108.33 | −12.1 | 3.65 | 1.1 | 11.76 |
| 0.0162 | 0 | −0.4 | 10.11 | −75.5 | 54.33 | −19.3 | 117.90 | −11.5 | 5.85 | 1.1 | 11.77 |
| 0.0207 | 0 | −0.6 | 10.15 | −79.0 | 70.58 | −22.6 | 119.98 | −8.8 | 7.31 | 0.8 | 11.47 |
| 0.0264 | 0 | −0.6 | 10.22 | −81.7 | 90.83 | −24.7 | 122.06 | −8.7 | 6.40 | 0.6 | 11.25 |
| 0.0336 | 0 | −0.6 | 10.26 | −83.9 | 115.44 | −27.9 | 125.14 | −6.5 | 4.09 | 0.8 | 11.15 |
| 0.0428 | 0 | −0.8 | 10.39 | −84.8 | 129.36 | −30.4 | 128.28 | −0.4 | 3.21 | 1.1 | 11.17 |
| 0.0546 | 0 | −0.9 | 10.47 | −85.3 | 156.56 | −32.5 | 130.51 | 0.0 | 3.21 | 1.2 | 11.21 |
| 0.0695 | 0 | −1.0 | 10.51 | −84.7 | 185.90 | −32.8 | 130.79 | 0.0 | 3.28 | 1.3 | 11.21 |
| 0.0886 | 0 | −1.2 | 10.58 | −86.7 | 206.53 | −32.8 | 130.71 | 0.0 | 3.35 | 1.5 | 11.24 |
| 0.1129 | 0 | −1.6 | 10.61 | −85.2 | 219.63 | −31.9 | 129.34 | 0.0 | 3.41 | 2.0 | 11.23 |
| 0.1438 | 0 | −2.0 | 10.57 | −86.0 | 225.62 | −31.5 | 129.14 | 0.0 | 3.44 | 2.7 | 11.18 |
| 0.1833 | 0 | −2.0 | 10.51 | −87.3 | 224.84 | −30.0 | 127.03 | −0.1 | 3.46 | 3.0 | 11.31 |
| 0.2336 | 0 | −2.0 | 10.48 | −85.3 | 220.11 | −30.2 | 126.17 | −0.1 | 3.47 | 2.4 | 11.42 |
| 0.2976 | 0 | −2.3 | 10.47 | −86.9 | 219.66 | −30.1 | 126.87 | −0.1 | 3.48 | −0.4 | 11.37 |
| 0.3793 | 0 | −2.6 | 10.47 | −85.3 | 214.48 | −30.3 | 126.06 | −0.1 | 3.49 | −4.3 | 11.28 |
| 0.4833 | 0 | −2.8 | 10.48 | −85.0 | 207.07 | −29.6 | 126.40 | −0.1 | 3.50 | −9.0 | 11.52 |
| 0.6158 | 0 | −2.8 | 10.53 | −86.5 | 198.52 | −29.4 | 125.98 | −0.1 | 3.51 | −11.5 | 11.85 |
| 0.7848 | 0 | −2.8 | 10.61 | −85.7 | 191.85 | −28.8 | 125.52 | −0.1 | 3.52 | −15.9 | 11.94 |
| 1.0000 | 0 | −3.0 | 10.64 | −83.4 | 186.43 | −25.9 | 121.61 | −0.1 | 3.52 | −23.8 | 11.77 |
| MAE ± SD | | 3.2 ± 7.2 | | 82.9 ± 5.1 | | 28.0 ± 4.3 | | 2.9 ± 4.4 | | 4.3 ± 6.0 | |

Table S5: Contrast saturation. Perceived motion strength should increase with contrast at low levels and saturate at higher levels; the table follows the response as contrast increases.

| Mask (c/°) | Direction (°) | RAFT $\hat{\theta}\pm$ SD (°) | mag | PWC-Net $\hat{\theta}\pm$ SD (°) | mag | LFN2 $\hat{\theta}\pm$ SD (°) | mag | ME-Att $\hat{\theta}\pm$ SD (°) | mag | Dual $\hat{\theta}\pm$ SD (°) | mag |
|---|---|---|---|---|---|---|---|---|---|---|---|
| no mask | 0 | $-2.8 \pm 15.2$ | 11.80 | $-77.1 \pm 1.3$ | 266.82 | $7.8 \pm 6.0$ | 13.47 | $1.7 \pm 0.0$ | 3.53 | $-28.3 \pm 2.0$ | 19.86 |
| 0.02 | 0 | $-31.6 \pm 25.7$ | 13.48 | $-78.2 \pm 2.1$ | 272.07 | $6.4 \pm 8.4$ | 14.84 | $1.9 \pm 0.1$ | 3.54 | $-154.4 \pm 42.5$ | 30.65 |
| 0.05 | 0 | $-83.0 \pm 25.3$ | 44.69 | $-77.5 \pm 1.9$ | 273.06 | $6.3 \pm 8.1$ | 14.41 | $2.1 \pm 0.2$ | 3.57 | $-170.9 \pm 22.5$ | 31.87 |
| 0.1 | 0 | $-94.6 \pm 45.3$ | 55.32 | $-78.4 \pm 1.6$ | 271.40 | $4.1 \pm 7.4$ | 13.70 | $2.4 \pm 0.3$ | 3.64 | $-177.6 \pm 5.0$ | 38.48 |
| 0.2 | 0 | $-105.1 \pm 80.8$ | 65.28 | $-78.4 \pm 2.0$ | 268.85 | $2.0 \pm 7.9$ | 14.11 | $5.0 \pm 1.1$ | 4.09 | $179.7 \pm 6.5$ | 46.11 |
| 0.25 | 0 | $148.7 \pm 110.5$ | 54.55 | $-78.6 \pm 2.4$ | 269.11 | $2.6 \pm 7.4$ | 14.25 | $5.8 \pm 1.5$ | 4.31 | $179.7 \pm 9.7$ | 44.40 |
| 0.5 | 0 | $-74.9 \pm 37.8$ | 63.36 | $-79.0 \pm 3.3$ | 262.75 | $-4.9 \pm 8.3$ | 15.31 | $14.9 \pm 30.1$ | 7.64 | $-172.2 \pm 48.8$ | 38.57 |
| 0.75 | 0 | $-95.3 \pm 10.7$ | 158.37 | $-79.2 \pm 3.6$ | 255.69 | $-10.3 \pm 12.4$ | 19.06 | $34.6 \pm 52.3$ | 8.28 | $-150.7 \pm 53.8$ | 35.65 |
| 1 | 0 | $134.0 \pm 96.9$ | 17.21 | $-78.9 \pm 2.9$ | 252.94 | $15.4 \pm 66.2$ | 8.08 | $18.3 \pm 60.3$ | 6.04 | $-147.3 \pm 61.4$ | 33.63 |
| 1.25 | 0 | $121.7 \pm 28.0$ | 65.87 | $-79.8 \pm 2.1$ | 239.83 | $47.3 \pm 93.5$ | 10.96 | $-3.1 \pm 44.8$ | 5.24 | $-167.4 \pm 63.6$ | 31.98 |
| 1.5 | 0 | $85.7 \pm 47.1$ | 29.36 | $-79.7 \pm 2.5$ | 237.37 | $13.1 \pm 70.9$ | 11.31 | $4.5 \pm 41.9$ | 5.38 | $-163.5 \pm 59.0$ | 31.51 |
| 2 | 0 | $-23.3 \pm 41.0$ | 40.07 | $-82.3 \pm 2.8$ | 243.82 | $-13.6 \pm 36.5$ | 10.72 | $23.3 \pm 30.7$ | 5.25 | $-169.9 \pm 43.4$ | 29.66 |
| 2.5 | 0 | $-147.5 \pm 56.1$ | 13.93 | $-80.0 \pm 2.4$ | 259.51 | $-19.5 \pm 34.4$ | 25.31 | $49.6 \pm 57.1$ | 5.41 | $-176.9 \pm 75.2$ | 22.08 |
| 3 | 0 | $126.2 \pm 118.3$ | 26.81 | $-81.0 \pm 1.5$ | 271.62 | $-3.2 \pm 16.2$ | 16.77 | $26.9 \pm 40.5$ | 2.29 | $173.6 \pm 55.8$ | 35.66 |
| 4 | 0 | $103.6 \pm 26.9$ | 48.36 | $-87.5 \pm 2.0$ | 284.09 | $157.8 \pm 93.1$ | 9.96 | $-176.0 \pm 68.0$ | 5.75 | $117.5 \pm 128.3$ | 25.91 |
| 5 | 0 | $-66.9 \pm 31.9$ | 47.49 | $-84.5 \pm 1.4$ | 319.50 | $18.4 \pm 12.3$ | 10.13 | $-61.1 \pm 108.7$ | 7.23 | $177.4 \pm 69.2$ | 20.47 |
| 6 | 0 | $85.9 \pm 8.7$ | 154.63 | $-85.6 \pm 2.0$ | 339.61 | $41.1 \pm 21.3$ | 5.09 | $10.3 \pm 13.5$ | 8.10 | $-162.9 \pm 64.6$ | 25.84 |
| 8 | 0 | $-92.0 \pm 35.9$ | 40.59 | $-77.3 \pm 0.3$ | 338.25 | $25.6 \pm 11.4$ | 8.35 | $-0.7 \pm 3.5$ | 7.84 | $125.7 \pm 71.7$ | 20.13 |
| 10 | 0 | $-71.5 \pm 53.5$ | 41.10 | $-77.0 \pm 2.0$ | 320.67 | $-11.9 \pm 12.0$ | 10.28 | $-7.5 \pm 6.6$ | 6.28 | $156.9 \pm 43.7$ | 24.00 |
| 12 | 0 | $-79.6 \pm 84.1$ | 16.88 | $-76.3 \pm 0.3$ | 333.82 | $42.2 \pm 48.7$ | 8.06 | $-25.2 \pm 15.8$ | 8.67 | $106.3 \pm 74.3$ | 32.24 |
| MAE ± SD | | $83.3 \pm 53.3$ | | $79.8 \pm 3.7$ | | $29.2 \pm 41.0$ | | $28.5 \pm 46.4$ | | $133.6 \pm 59.1$ | |

Table S6: Motion masking. The first column represents the spatial frequency of the mask, and "no mask" represents the signal presented alone. The ground-truth direction is 0° in each row. The values are averaged across 50 presentations with varying spatial phases of the stimuli.

| Stimulus | Direction (°) | RAFT $\hat{\theta}$ (°) | mag | PWC-Net $\hat{\theta}$ (°) | mag | LFN2 $\hat{\theta}$ (°) | mag | ME-Att $\hat{\theta}$ (°) | mag | Dual $\hat{\theta}$ (°) | mag |
|---|---|---|---|---|---|---|---|---|---|---|---|
| Square wave | 0 | 1.8 | 24.13 | −26.6 | 26.96 | −32.7 | 28.41 | 2.7 | 24.17 | 5.6 | 22.24 |
| Missing fund. | 180 | 2.6 | 24.05 | −82.7 | 326.46 | −38.8 | 30.72 | 174.7 | 11.13 | 1.5 | 27.12 |
| MAE ± SD | | $89.6 \pm 87.8$ | | $62.0 \pm 35.4$ | | $87.0 \pm 54.2$ | | $4.0 \pm 1.3$ | | $92.1 \pm 86.4$ | |

Table S7: Missing-fundamental illusion.

| Stimulus | Direction (°) | RAFT $\hat{\theta}\pm$ SD (°) | mag | PWC-Net $\hat{\theta}\pm$ SD (°) | mag | LFN2 $\hat{\theta}\pm$ SD (°) | mag | ME-Att $\hat{\theta}\pm$ SD (°) | mag | Dual $\hat{\theta}\pm$ SD (°) | mag |
|---|---|---|---|---|---|---|---|---|---|---|---|
| Rightward | 0 | $99.0 \pm 22.0$ | 143.64 | $-87.6 \pm 2.2$ | 249.95 | $-80.4 \pm 8.0$ | 48.93 | $6.2 \pm 5.9$ | 2.15 | $-11.5 \pm 23.4$ | 20.70 |
| Leftward | 180 | $91.4 \pm 19.6$ | 126.28 | $-91.1 \pm 1.9$ | 248.08 | $-112.2 \pm 92.3$ | 19.41 | $179.4 \pm 0.5$ | 2.43 | $-176.0 \pm 7.6$ | 19.87 |
| MAE ± SD | | $94.2 \pm 21.7$ | | $88.3 \pm 2.1$ | | $82.2 \pm 30.7$ | | $4.0 \pm 4.5$ | | $13.9 \pm 13.3$ | |

Table S8: Second-order motion. Values are averaged across the 100 presentations with different random noise carriers for each direction.

| Stimulus | Direction (°) | RAFT $\hat{\theta}\pm$ SD (°) | mag | PWC-Net $\hat{\theta}\pm$ SD (°) | mag | LFN2 $\hat{\theta}\pm$ SD (°) | mag | ME-Att $\hat{\theta}\pm$ SD (°) | mag | Dual $\hat{\theta}\pm$ SD (°) | mag |
|---|---|---|---|---|---|---|---|---|---|---|---|
| Phi | 0 | $75.6 \pm 22.2$ | 91.84 | $-25.0 \pm 45.4$ | 20.00 | $-55.1 \pm 14.9$ | 22.22 | $-18.4 \pm 11.1$ | 7.21 | $-18.1 \pm 8.1$ | 12.05 |
| Reverse-Phi | 180 | $101.9 \pm 19.2$ | 153.50 | $-91.8 \pm 18.0$ | 102.35 | $-77.1 \pm 33.5$ | 62.66 | $179.3 \pm 2.0$ | 1.69 | $-178.4 \pm 38.7$ | 43.26 |
| MAE ± SD | | $76.1 \pm 21.1$ | | $65.3 \pm 31.2$ | | $78.9 \pm 35.3$ | | $9.8 \pm 11.7$ | | $20.0 \pm 35.3$ | |

Table S9: Phi and reverse-Phi illusions. Values are averaged across 20 presentations with a different random noise pattern each.

| Model | Forward | | | Reversed | | | $V/I$ |
|---|---|---|---|---|---|---|---|
| | Direction (°) | $\hat{\theta}\pm$ SD (°) | mag | Direction (°) | $\hat{\theta}\pm$ SD (°) | mag | |
| *Light edges* | | | | | | | |
| RAFT | 0 | $50.6 \pm 28.5$ | 7.94 | 180 | $107.2 \pm 49.8$ | 4.65 | 1.88 |
| PWC-Net | 0 | $-45.3 \pm 20.3$ | 10.40 | 180 | $-114.4 \pm 4.3$ | 16.85 | $-20.42$ |
| LFN2 | 0 | $-51.5 \pm 3.8$ | 8.81 | 180 | $90.7 \pm 19.9$ | 7.10 | 0.98 |
| ME-Att | 0 | $-2.9 \pm 0.2$ | 2.92 | 180 | $-179.8 \pm 0.0$ | 3.21 | $-21.21$ |
| Dual | 0 | $15.1 \pm 8.8$ | 4.02 | 180 | $-170.0 \pm 12.8$ | 2.51 | 4.36 |
| *Uneven square wave* | | | | | | | |
| RAFT | 0 | $15.8 \pm 15.3$ | 11.02 | 180 | $178.8 \pm 0.2$ | 9.38 | 28.49 |
| PWC-Net | 0 | $8.6 \pm 1.4$ | 10.09 | 180 | $132.9 \pm 0.7$ | 14.57 | 361.07 |
| LFN2 | 0 | $-69.4 \pm 0.4$ | 28.44 | 180 | $-148.4 \pm 4.0$ | 11.97 | $-122.88$ |
| ME-Att | 0 | $-29.7 \pm 1.0$ | 11.45 | 180 | $-179.1 \pm 0.2$ | 6.81 | 5.35 |
| Dual | 0 | $-12.5 \pm 1.9$ | 11.58 | 180 | $175.6 \pm 1.3$ | 9.94 | 15.34 |
| *Opponency* (control) | | | | | | | |
| Bright bar | | | | | | | |
| RAFT | 0 | 54.3 | 19.08 | 180 | 173.6 | 11.06 | 155.93 |
| PWC-Net | 0 | $-57.2$ | 19.73 | 180 | $-114.8$ | 24.10 | 35.82 |
| LFN2 | 0 | $-56.6$ | 24.00 | 180 | 110.8 | 23.90 | 4.59 |
| ME-Att | 0 | $-2.9$ | 7.47 | 180 | $-179.9$ | 5.74 | 7.67 |
| Dual | 0 | 25.2 | 2.28 | 180 | $-179.9$ | 2.17 | $-39.56$ |
| Dark bar | | | | | | | |
| RAFT | 0 | 13.6 | 10.44 | 180 | 156.9 | 4.45 | 2.35 |
| PWC-Net | 0 | $-60.6$ | 24.19 | 180 | $-113.9$ | 29.60 | $-204.34$ |
| LFN2 | 0 | $-60.2$ | 16.21 | 180 | 171.5 | 8.33 | $-89.24$ |
| ME-Att | 0 | $-1.0$ | 5.29 | 180 | $-179.9$ | 4.06 | 7.61 |
| Dual | 0 | 11.2 | 2.76 | 180 | $-173.3$ | 2.25 | 10.45 |

Table S10: Breaking of time-reversal symmetry. $V/I$ measures how much of the response breaks the symmetry. For light edges and the uneven square wave, values are averaged across 20 spatial phases. The bright and dark moving bars are the control, for which no symmetry breaking is expected.

### Supplementary results of the proposed model

The following figures show the responses of the proposed model to the stimuli described in the Methods section.

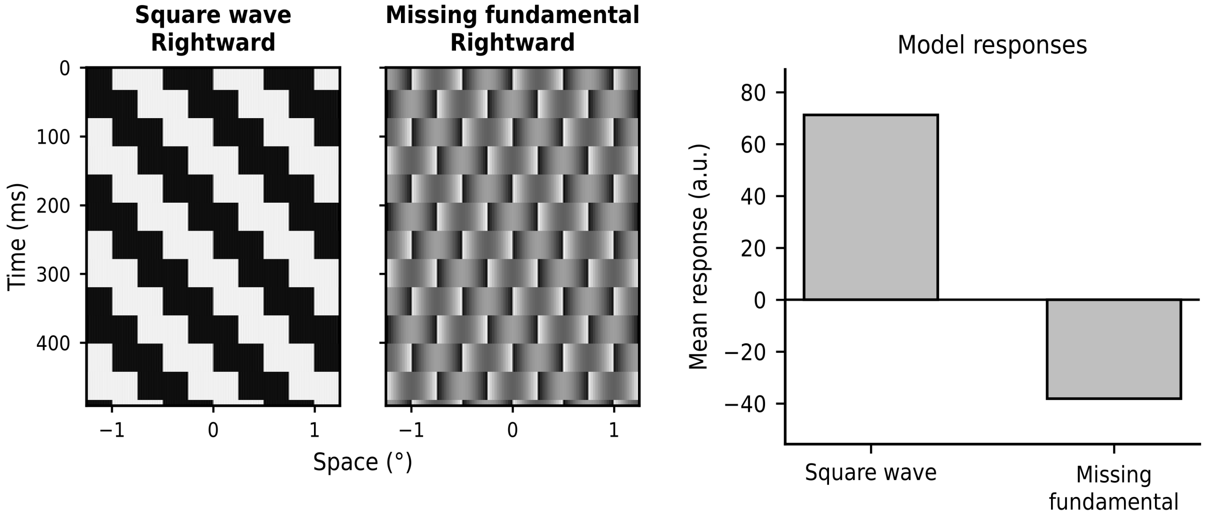


Figure S1: The proposed model reproduces the missing-fundamental illusion. Left: Space-time plots of a square-wave grating moving rightward in discrete steps and of the same grating after removing its fundamental Fourier component. Right: The model's response to the square wave is positive, correctly indicating rightward motion, whereas its response to the missing-fundamental stimulus is negative, indicating leftward motion.

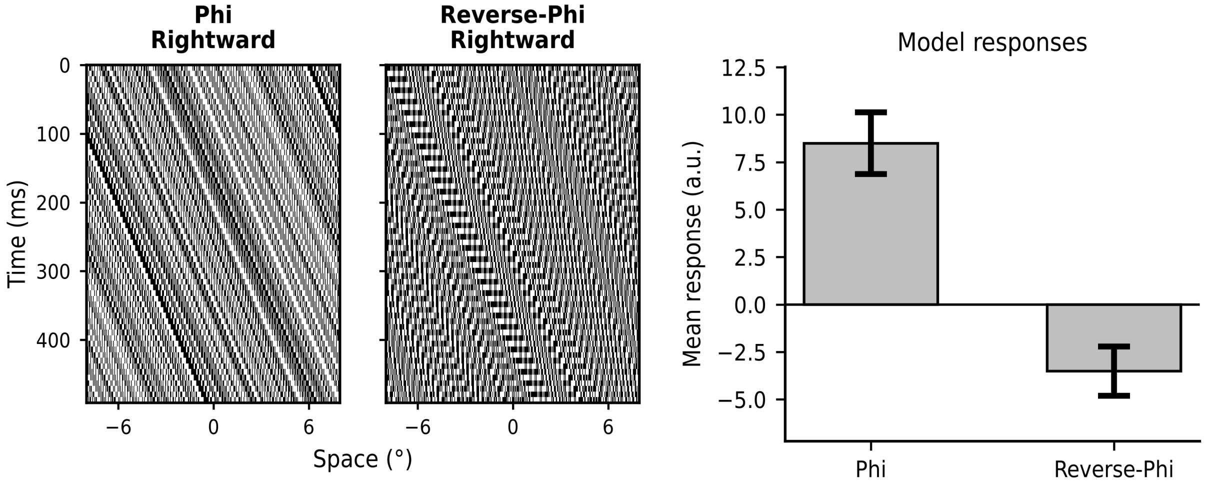


Figure S2: The proposed model reproduces the reverse-Phi illusion. Left: Space-time plots of a random contrast pattern moving rightward, which elicits ordinary Phi motion, and of the same sequence with its contrast polarity inverted at every frame, which elicits reverse-Phi motion. Right: The model's response is positive for the Phi stimulus, correctly indicating rightward motion, and negative for the reverse-Phi stimulus, indicating leftward motion. Responses are averaged across different random generations of the stimuli, and error bars show the SD.

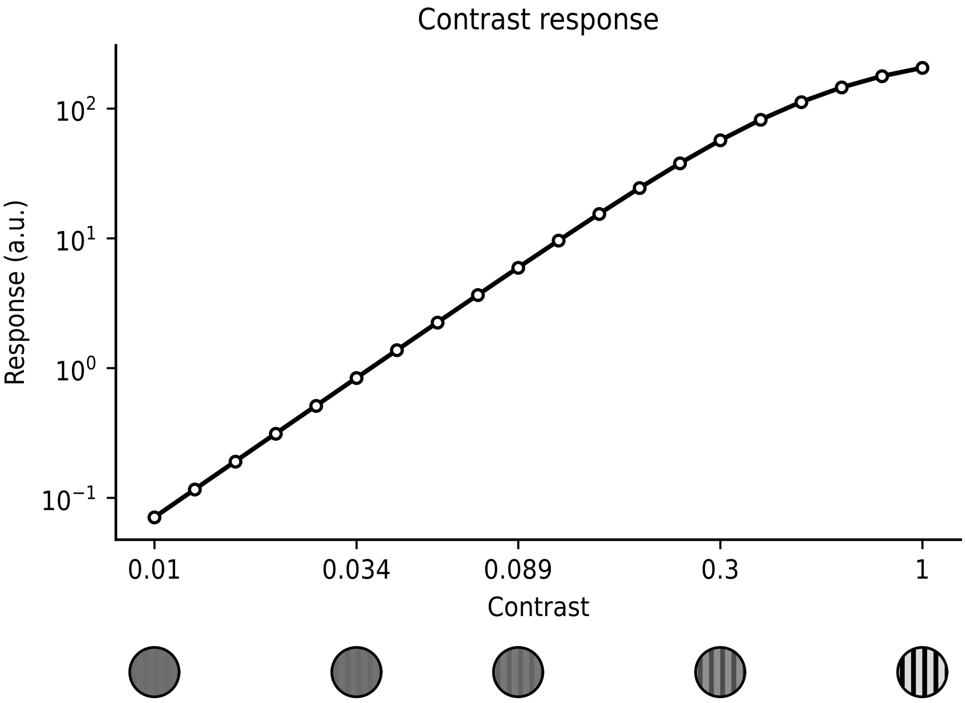


Figure S3: The model reproduces contrast saturation. The magnitude of the model's response to a drifting sinusoidal grating as a function of its contrast, shown on a log–log scale; the icons illustrate the grating at some of the contrasts tested. The response grows monotonically with contrast at low contrast levels but saturates at higher contrasts.

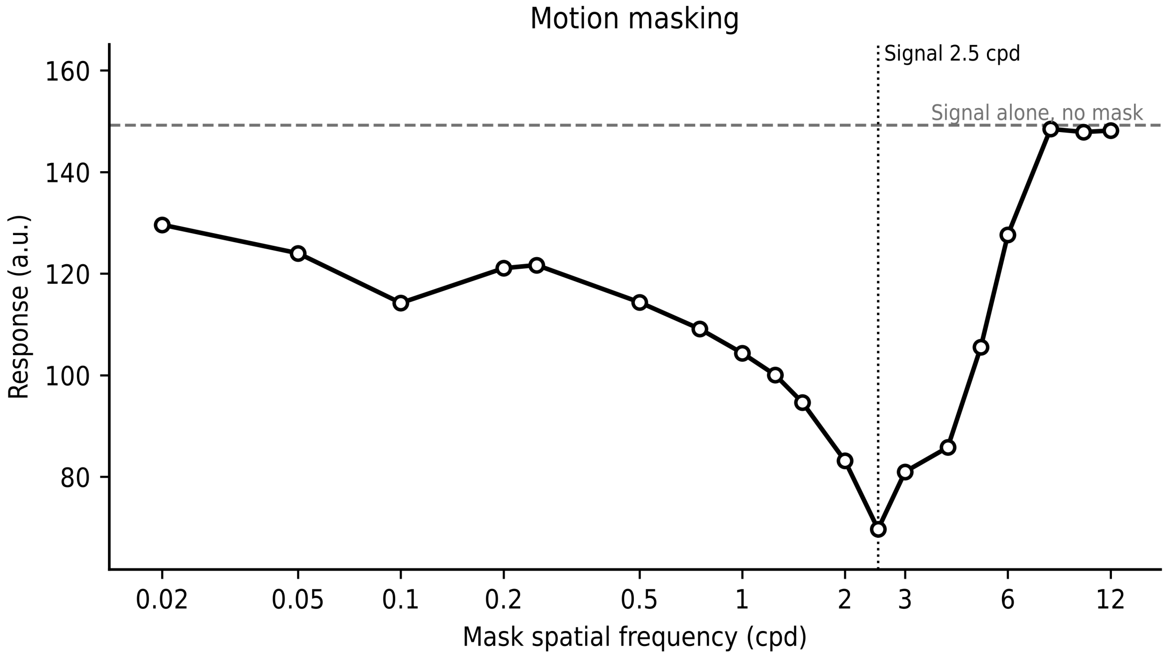


Figure S4: The proposed model reproduces motion masking. The magnitude of the model's response to a signal grating presented together with a jittering mask grating, as a function of the mask's spatial frequency; the dashed line marks the response to the signal alone, and the dotted line marks the spatial frequency of the signal itself. The response is suppressed the most when the mask and the signal share a similar spatial frequency, and recovers as the mask's frequency moves away from that of the signal, reproducing the frequency-tuned nature of motion masking. Responses are averaged across 50 presentations with varying spatial phase of the stimuli.

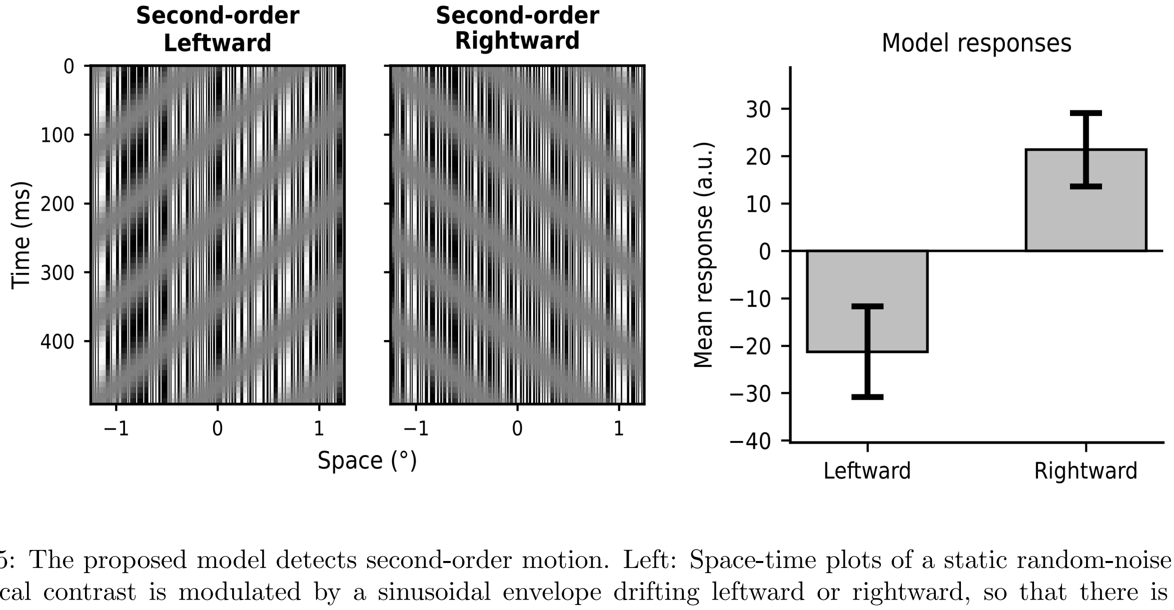


Figure S5: The proposed model detects second-order motion. Left: Space-time plots of a static random-noise carrier whose local contrast is modulated by a sinusoidal envelope drifting leftward or rightward, so that there is no net displacement of mean luminance. Right: The model's response is negative for the leftward stimulus and positive for the rightward stimulus, correctly recovering the direction of motion of the envelope. Responses are averaged across 100 presentations with a different random noise carrier each, and error bars show the SD.

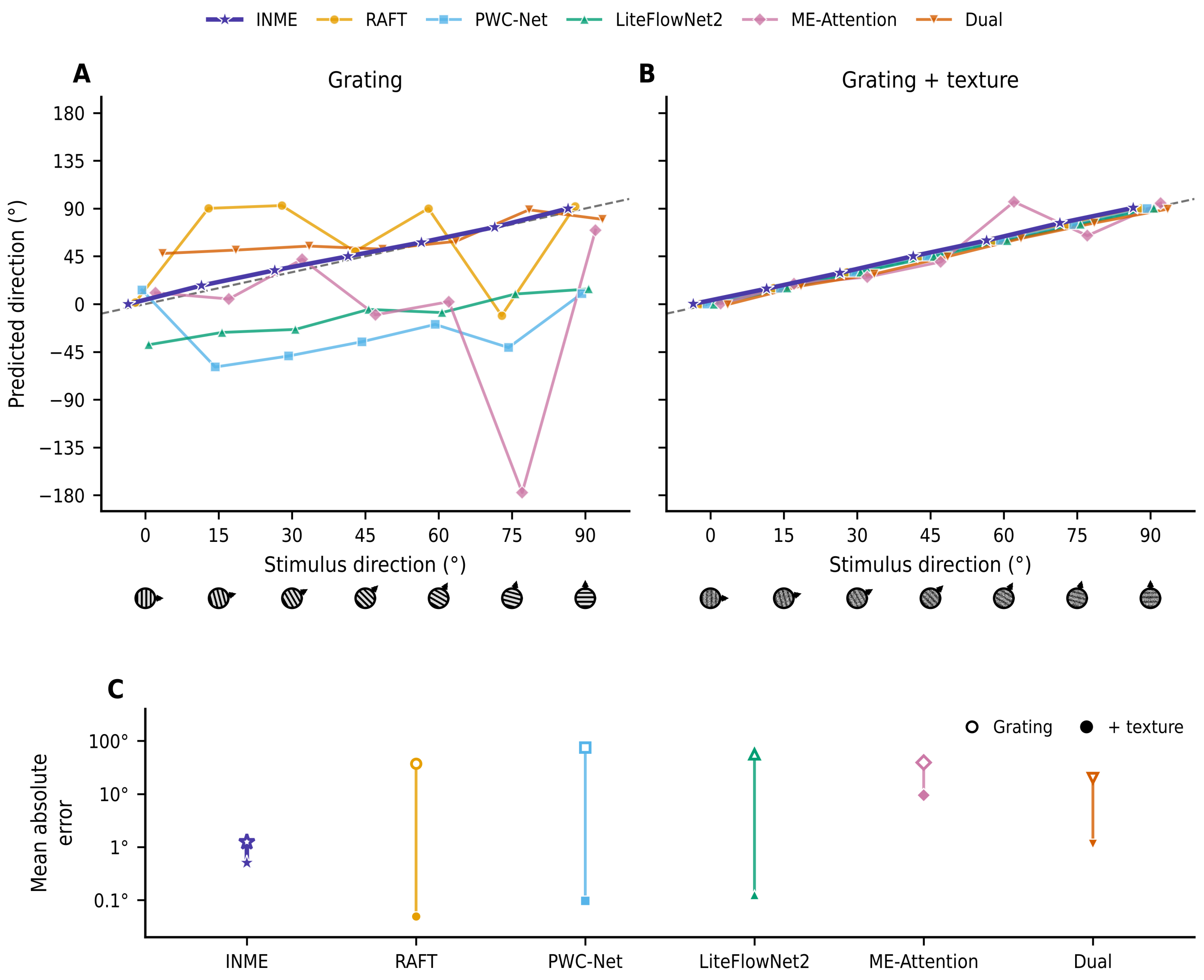


Figure S6: Adding texture to the grating markedly improves the DNNs' direction estimates, whereas the proposed model (INME) is accurate with or without it. (A) Predicted versus stimulus direction for a drifting grating; the dashed line marks perfect agreement, and the icons below the axis show the stimulus at each direction. (B) The same, with a moving band-pass noise texture added on top of the grating. (C) Mean absolute error of each model in the two conditions on a logarithmic scale: open symbols for the grating alone, and filled symbols with the texture added.
